# Specialization of the *Drosophila* nuclear export family protein, Nxf3, for piRNA precursor export

**DOI:** 10.1101/716258

**Authors:** Emma Kneuss, Marzia Munafò, Evelyn L. Eastwood, Undine-Sophie Deumer, Jonathan B. Preall, Gregory J. Hannon, Benjamin Czech

**Affiliations:** Cancer Research UK Cambridge Institute, University of Cambridge, Li Ka Shing Centre, Cambridge CB2 0RE, United Kingdom; Cold Spring Harbor Laboratory, Cold Spring Harbor, NY 11724, USA

**Keywords:** transposon control, PIWI proteins, piRNA clusters, nuclear export factor, RNA export

## Abstract

The piRNA pathway is a conserved, small RNA-based immune system that protects animal germ cell genomes from the harmful effects of transposon mobilisation. In *Drosophila* ovaries, most piRNAs originate from dual-strand clusters, which generate piRNAs from both genomic strands. Dual-strand clusters use non-canonical transcription mechanisms. Although transcribed by RNA polymerase II, cluster transcripts lack splicing signatures and poly(A) tails. mRNA processing is important for general mRNA export mediated by Nuclear export factor 1. Although UAP56, a component of the transcription and export complex, has been implicated in piRNA precursor export, it remains unknown how dual-strand cluster transcripts are specifically targeted for piRNA biogenesis by export from the nucleus to cytoplasmic processing centers. Here we report that dual-strand cluster transcript export requires CG13741/Bootlegger and the *Drosophila* Nuclear export factor family protein, Nxf3. Bootlegger is specifically recruited to piRNA clusters and in turn brings Nxf3. We find that Nxf3 specifically binds to piRNA precursors and is essential for their export to piRNA biogenesis sites, a process that is critical for germline transposon silencing. Our data shed light on how dual-strand clusters compensate for a lack of canonical features of mature mRNAs to be specifically exported via Nxf3, ensuring proper piRNA production.

## Introduction

Transposable elements represent a major threat to genome integrity. In the animal germline, transposon mobilization is prevented by a conserved small RNA-based immune system, the PIWI-interacting RNA (piRNA) pathway. piRNAs are 23-to 30-nucleotide (nt) small RNAs that associate with Argonaute proteins of the PIWI clade (Czech et al. 2018; Ozata et al. 2019). The majority of piRNAs are produced from discrete genomic loci, called piRNA clusters, which are composed of transposon remnants, and thus constitute a genetic memory of past transposon exposure (Brennecke et al. 2007). In *Drosophila*, piRNA clusters give rise to piRNA precursor transcripts from either one (uni-strand clusters) or both (dual-strand clusters) genomic strands (Malone et al. 2009; Mohn et al. 2014). The processing of precursor transcripts into mature piRNAs is a cytoplasmic event, occurring in nuage, cytoplasmic structures adjacent to the nucleus which are enriched for piRNA biogenesis components (Lim and Kai 2007; Malone et al. 2009). Precursor RNAs, therefore, require export from the nucleus for conversion into small RNAs. Yet, how precursors are identified for export and selectively trafficked to nuage remains a mystery.

Canonical mRNA export in eukaryotes relies on Nuclear export factor 1 (Nxf1) and its cofactor Nxt1 (NTF2-related export protein 1) (Herold et al. 2001; Kohler and Hurt 2007; Stewart 2010). However, mRNA export is a highly regulated process, occurring only on transcripts that have undergone mRNA processing. This includes the formation of the 5’ cap, splicing, and 3’ end processing, which involves endo-nucleolytic cleavage followed by polyadenylation (Le Hir et al. 2001; Cheng et al. 2006; Kohler and Hurt 2007; Yoh et al. 2007). Completion of mRNA processing leads to the recruitment of the transcription and export complex (TREX), which is a prerequisite for association with the Nxf1-Nxt1 heterodimer. Uni-strand clusters, such as *flamenco* (*flam*), are capped, spliced, and polyadenylated, and consequently follow canonical export rules, relying on Nxf1-Nxt1 (Goriaux et al. 2014; Mohn et al. 2014; Dennis et al. 2016).

In contrast, germline-specific dual-strand clusters produce non-canonical transcripts that, while capped and transcribed by RNA polymerase II (Pol II), lack splicing signatures and poly(A) tails (Mohn et al. 2014; Zhang et al. 2014; Andersen et al. 2017). Furthermore, H3K4me2 patterns typical of active promoters are missing on dual-strand clusters. Instead, these loci are embedded within heterochromatin and feature H3K9me3 marks, which are normally not correlated with transcription (Mohn et al. 2014; Zhang et al. 2014). Recent work has uncovered the machinery that facilitates heterochromatin-dependent dual-strand cluster transcription (Mohn et al. 2014; Zhang et al. 2014; Andersen et al. 2017). The Heterochromatin Protein 1a (HP1a) homolog Rhino (Rhi) specifically associates with dual-strand clusters through its H3K9me3 binding domain (Klattenhoff et al. 2009; Mohn et al. 2014; Zhang et al. 2014). Rhi interacts with the linker protein Deadlock (Del), which was shown to recruit Moonshiner (Moon), a homologue of the transcription initiation factor II A (TFIIA) (Andersen et al. 2017). Moon promotes transcription initiation of dual-strand cluster loci via its interaction with the TATA-binding protein (TBP)-related factor TRF2. Del also interacts with Cutoff (Cuff), a protein with similarity to the Rai1 transcription termination factor (Pane et al. 2011; Mohn et al. 2014; Zhang et al. 2014; Chen et al. 2016). Cuff suppresses splicing and Pol II termination and is thought to protect dual-strand piRNA cluster transcripts from degradation by the exonuclease Rat1 (Zhang et al. 2014; Chen et al. 2016). Thus, this Rhi-anchored complex generates transcripts that lack signatures of canonical mRNAs. UAP56, a DEAD-box helicase also involved in canonical mRNA splicing and export, was shown to associate with piRNA precursor transcripts (Zhang et al. 2012; Zhang et al. 2018). Co-localization of UAP56 with the sites of dual-strand cluster transcription and a close association with the perinuclear DEAD-box helicase Vasa (Vas) suggested that this protein is involved in coupling precursor synthesis to downstream processing (Zhang et al. 2012).

The non-canonical transcription of dual-strand clusters (absence of splicing and polyadenylation) suggests that their RNA products require an alternative export machinery that does not involve Nxf1/Nxt1. Here, we show that piRNA cluster export depends on a germline-specific paralogue of Nxf1, Nuclear export factor 3 (Nxf3), that is recruited to cluster transcripts through its interaction with CG13741/Bootlegger and the Rhino-Deadlock-Cutoff (RDC) complex. Both CG13741/Bootlegger and Nxf3 are essential for piRNA-guided transposon repression in germ cells. Nxf3-dependent export of piRNA precursors from the nucleus to nuage likely involves Nxt1, and uses a Crm1 (Chromosomal maintenance 1)-dependent mechanism. Together, our data shed light on how non-canonical piRNA precursor transcripts are exported from their heterochromatic source loci to the processing sites in the cytoplasm and how specific components of the nuclear export machinery are adapted to facilitate this.

## Results

### CG13741 and Nxf3 are germline-specific piRNA factors involved in dual-strand cluster biology

Genome-wide screens in *Drosophila* have revealed the genetic framework of the piRNA pathway in somatic and germ cells (Czech et al. 2013; Handler et al. 2013; Muerdter et al. 2013). Since proteins involved in dual-strand piRNA cluster biology will specifically compromise transposon silencing in germ cells, we focused on two uncharacterized germline-specific screen hits: CG13741 and Nuclear export factor 3 (Nxf3) (Czech et al. 2013). CG13741 is a 42 kDa protein without identifiable domains (**Fig. 1A**) that is expressed predominantly in ovaries (**Fig. S1A**). Nxf3 is also expressed preferentially in ovaries (**Fig. S1A**) and is a member of the nuclear export factor family, which in *Drosophila* includes three additional proteins (**Fig. S1B**). Ubiquitously expressed Nxf1 is responsible for bulk mRNA export (Herold et al. 2001), while ovary-enriched Nxf2 functions in piRNA-guided transcriptional transposon silencing (Batki et al. 2019; Fabry et al. 2019; Murano et al. 2019; Zhao et al. 2019). Nxf3, and the testis-specific Nxf4, have not yet been ascribed functions. Similar to other members of this protein family, Nxf3 contains an RNA-binding domain (RBD), leucin rich repeats (LRR), a NTF2-like domain (NTF2), and a diverged Ubiquitin-associated domain (UBA) (**Fig. 1B**). While Nxf1 and Nxf2 were identified to be required for transposon silencing in somatic and germline cells, Nxf3 was only required in germ cells (Czech et al. 2013; Handler et al. 2013; Muerdter et al. 2013).

**Figure 1.**
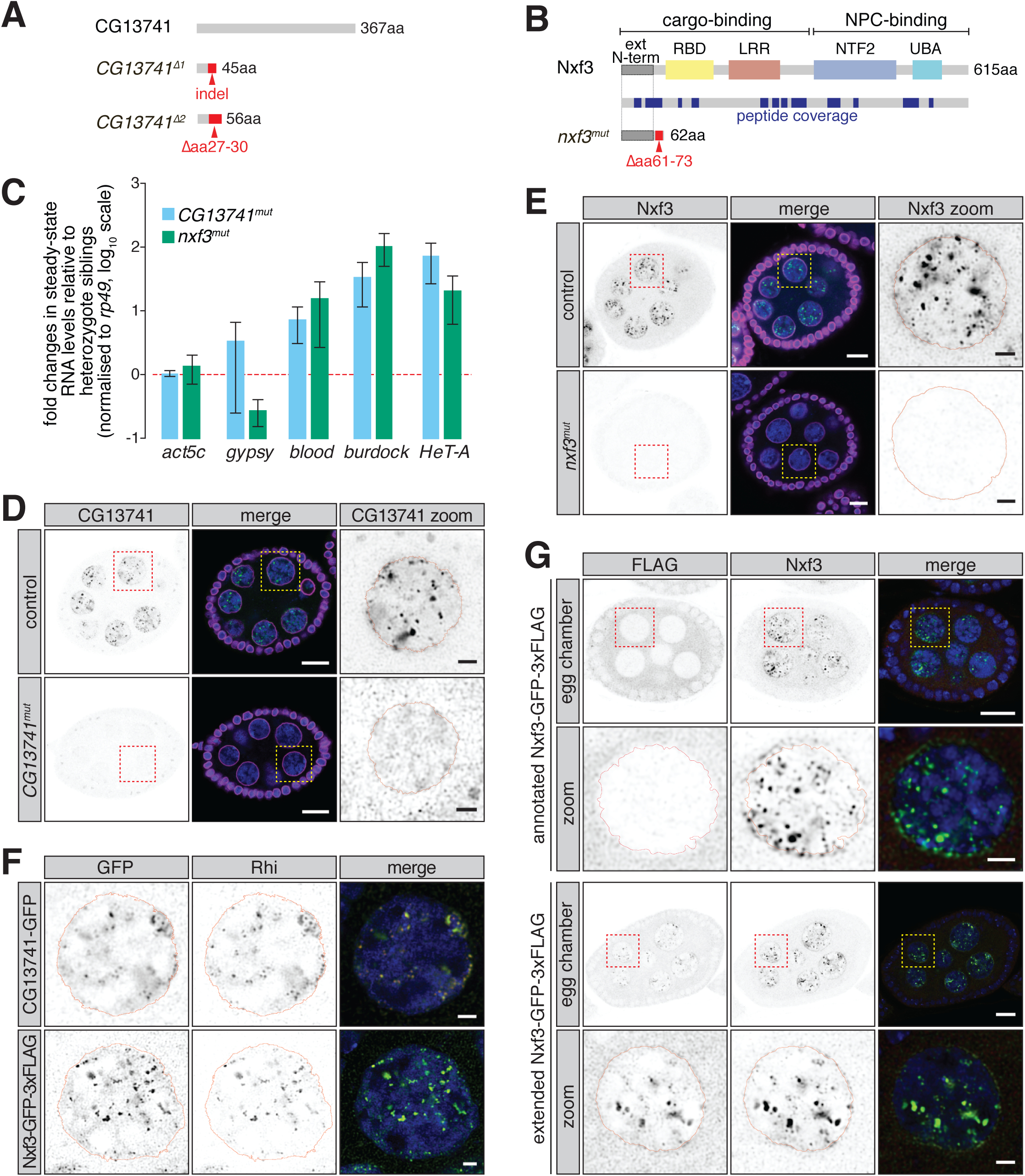
CG13741 and Nxf3 impact dual-strand piRNA cluster biology. **(A)** Schematic representation of CG13741, as well as *CG13741*^*Δ1*^ and *CG13741*^*Δ2*^ mutants. **(B)** Cartoon displaying the Nxf3 domain structure and the generated *nxf3*^*mut*^ allele. LRR, leucine rich repeats; RBD, RNA-binding domain; NTF2, NTF2-like domain; UBA, Ubiquitin associated domain. Peptide coverage obtained by Nxf3 pull-down followed by mass spectrometry is indicated in blue. **(C)** Bar graphs showing fold changes in steady-state RNA levels of a housekeeping gene (*act5c*), soma-specific (*gypsy*), intermediate (*blood*), and germline-specific (*HeT-A, burdock*) transposons in total ovarian RNA from the indicated genotypes (relative to heterozygote siblings and normalized to *rp49*). Error bars indicate standard deviation (n = 3). **(D)** Expression and localization of CG13741 in an egg chamber of control and *CG13741*^*mut*^ mutant flies is shown by immunofluorescence. A zoomed-in view of the indicated nurse cell nucleus is shown to the right. Green, CG13741; magenta, Lamin; blue, DNA. Scale bar for egg chamber, 10 µm; scale bar for zoom-in, 2 µm. **(E)** As in (D) but showing the expression and localization of Nxf3 in control and *nxf3*^*mut*^ egg chambers. Green, Nxf3; magenta, Lamin; blue, DNA. Scale bar for egg chamber, 10 µm; scale bar for zoom-in, 2 µm. **(F)** As in (D) but showing expression and localization of GFP-tagged CG13741 and GFP-tagged Nxf3 co-stained with Rhi. Green, GFP; red, Rhi; blue, DNA. Scale bar, 2 µm. **(G)** As in (D) but showing expression and localization of the annotated and extended Nxf3 sequence tagged C-terminally with GFP-3xFLAG and co-stained with an antibody to endogenous Nxf3. Green, Nxf3; red, FLAG; blue, DNA. Scale bar for egg chamber, 10 µm; scale bar for zoom-in, 2 µm.

Using CRISPR/Cas9, we generated *CG13741* and *nxf3* mutant flies. We obtained two *CG13741* mutant alleles, *CG13741*^*Δ1*^ and *CG13741*^*Δ2*^, which harbour premature stop codons that disrupt the CG13741 open reading frame from amino acid 27 onwards (**Fig. 1A**). For better readability, we refer to *CG13741* mutant and trans-heterozygote alleles as *CG13741*^*mut*^. For Nxf3, we recovered one mutant allele, *nxf3*^*mut*^, which leads to a premature stop codon that disrupts the open reading frame just downstream of the annotated start codon (**Fig. 1B**). Western blots on ovarian lysates from control, heterozygote and *nxf3* homozygous mutant flies confirmed this allele as null (**Fig. S1C** and see below). *CG13741* and *nxf3* mutants were female sterile, with homozygous mutant females laying slightly fewer eggs (∼70% of control females for both mutants), of which few (1.9% for *CG13741*^*mut*^) or none (0.0% for *nxf3*^*mut*^) hatched (**Fig. S1D**). Flies mutant for either *CG13741* or *nxf3* were compromised in the repression of intermediate (*blood*) and germline-specific transposons (*burdock, HeT-A*), while soma-specific transposable elements (*gypsy*) remained largely unchanged (**Fig. 1C**). We did not detect changes in the localization of piRNA-binding Argonaute proteins, Piwi, Aub or Ago3, in *CG13741*^*mut*^ and *nxf3*^*mut*^ ovaries (**Fig. S1E**). Thus, our data suggest that CG13741 and Nxf3 are important for transposon repression and could function in germline piRNA cluster biology.

In order to investigate their function in transposon control, we examined the expression pattern and subcellular localization of CG13741 and Nxf3 using polyclonal antibodies that we generated. CG13741 and Nxf3 both localized to discrete foci in nurse cell nuclei (**Fig. 1D,E**), which is highly reminiscent of staining patterns observed for RDC complex subunits (Klattenhoff et al. 2009; Pane et al. 2011; Mohn et al. 2014), while follicle cells lacked both proteins. CG13741 and Nxf3 both colocalise with nuclear foci stained by Rhi (**Fig. 1F**), supporting the hypothesis that they are involved in dual-strand piRNA cluster biology. Unlike RDC complex components, a fraction of Nxf3 (and to a lesser extent CG13741) also localizes to perinuclear foci that form ring-like structures outside nuclei and strongly resemble nuage (**Fig. 1D,E**), suggesting a possible role in piRNA precursor export.

To facilitate further biochemical studies, we created a Nxf3 cDNA transgene with a C-terminal epitope tag (GFP-3xFLAG). In contrast to the staining observed for the endogenous protein, the Nxf3 transgene was aberrantly localized to the cytoplasm (**Fig. 1G**), prompting us to examine the Nxf3 genomic locus. We noticed the presence of an additional 56 in-frame amino acids with an initiator methionine at the amino terminus. Immunoprecipitation of Nxf3 with our antibody followed by Mass Spectrometry (MS) revealed peptides specific to and overlapping with the putative N-terminal extension (**Fig. 1B**). We generated flies expressing the N-terminally extended Nxf3 sequence tagged carboxy-terminally with GFP-3xFLAG and found that the extended version colocalized with endogenous Nxf3 in germ cells as shown by immunofluorescence staining (**Fig. 1G**). We therefore conclude that the extended form of Nxf3 likely represents the functional and full-length protein. Unless specified otherwise, we used the extended version of Nxf3 for subsequent experiments and refer to this fusion as Nxf3-GFP-3xFLAG.

### CG13741 and Nxf3 recruitment to dual-strand clusters depends on the RDC complex

The RDC complex has an essential role in dual-strand cluster expression as it licenses clusters and recruits the transcription machinery to these loci (Mohn et al. 2014; Zhang et al. 2014). To understand the functional hierarchy of CG13741, Nxf3, and other piRNA pathway components, we depleted factors implicated in piRNA cluster biology in germ cells and examined the localization of CG13741, Nxf3, and other piRNA pathway components by immunofluorescence. Germline knockdown of all factors led to robust de-repression of intermediate and germline-specific transposons, indicating robust depletion (**Fig. S2A**). We first analyzed the dependency of Rhi and CG13741 on other known piRNA factors affecting dual-strand clusters by depleting these specifically in the germ cells of the ovary (**Fig. 2A**). Nuclear foci stained by Rhi were lost in germline knockdowns of components of the RDC complex, as reported previously (Mohn et al. 2014; Zhang et al. 2014). Depletion of Moon had no effect on Rhi recruitment, consistent with its function downstream of RDC (Andersen et al. 2017). Similarly, germline knockdown of *CG13741* and *nxf3* had no effect on Rhi localisation, suggesting that both factors function downstream of the RDC (**Fig. 2A**).

**Figure 2.**
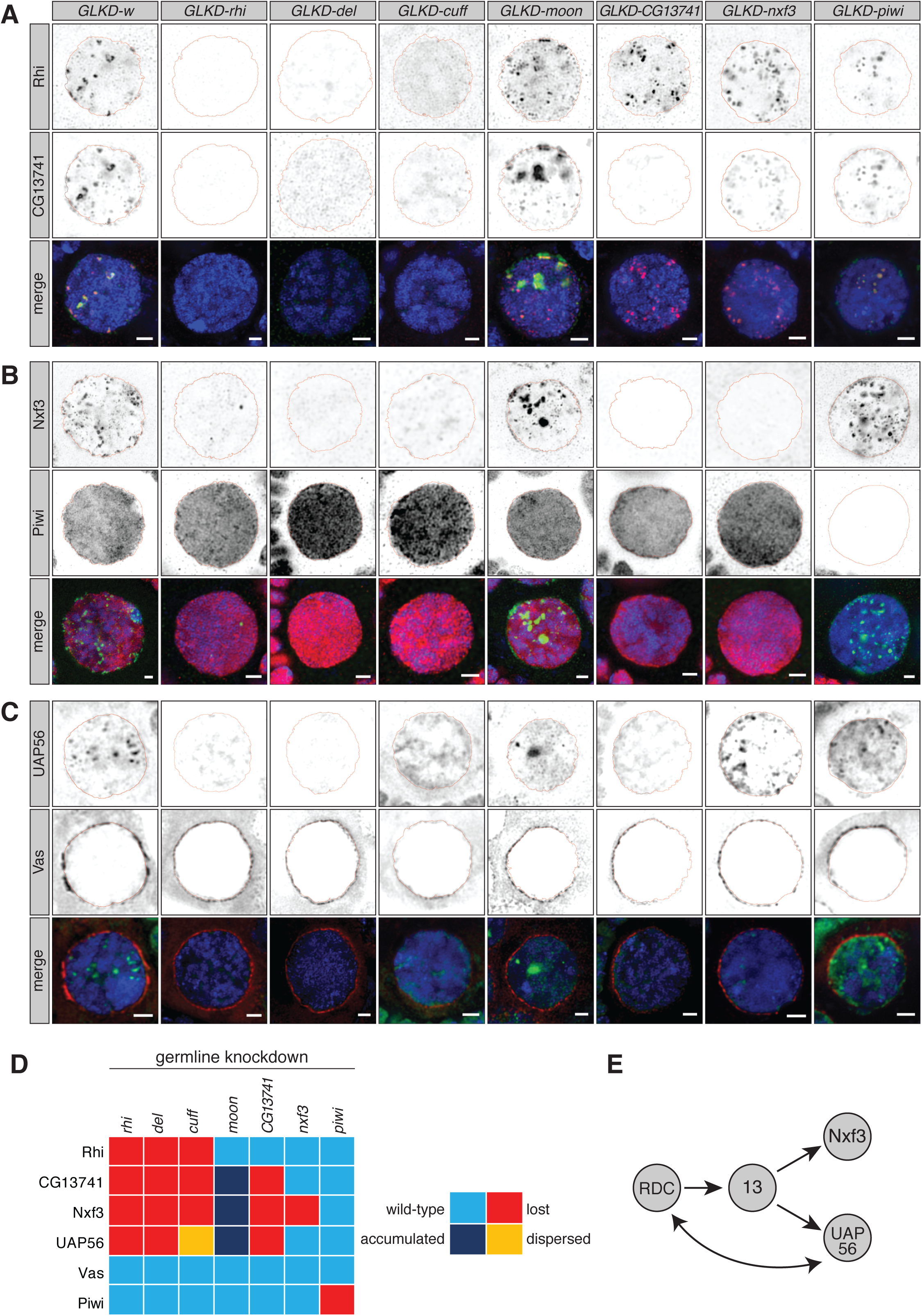
Hierarchy of dual-strand piRNA cluster factors. **(A)** Expression and localization of Rhi and CG13741 in nurse cell nuclei upon germline-specific knockdown (GLKD) of the indicated factors are shown by immunofluorescence. Green, CG13741; red, Rhi; blue, DNA. Scale bar, 2 µm. **(B)** As in (A) but showing the expression and localization of Piwi and Nxf3. Green, Nxf3; red, Piwi; blue, DNA. Scale bar, 2 µm. **(C)** As in (A) but showing the expression and localization of UAP56 and Vas. Green, UAP56; red, Vas; blue, DNA. Scale bar, 2 µm. **(D)** Heatmap summarizing the data shown in panels (A) through (C). **(E)** Cartoon showing the hierarchy of RDC, CG13741, UAP56, and Nxf3.

In control knockdowns, CG13741 was detected in nuclear foci that overlap with Rhi and a weak signal was detected outside nuclei (**Fig. 2A**). Recruitment of CG13741 to nuclear foci was dependent on all RDC complex components and completely lost upon germline-specific depletion of CG13741 itself (**Fig. 2A**). The localization of CG13741 to nuclear foci was unaffected by depletion of Nxf3. Interestingly, CG13741 signal outside of the nucleus was lost upon *nxf3* knockdown, suggesting a role of Nxf3 in the export of CG13741 out of the nucleus (**Fig. 2A**).

In control knockdowns, Nxf3 localized to discrete foci in nurse cell nuclei, likely piRNA cluster loci, as well as to nuage (**Fig. 2B**). Upon depletion of RDC complex components, all discrete concentrations of Nxf3 signal were lost. Additionally, the localisation of Nxf3 to nuclear foci was dependent on the presence of CG13741 (**Fig. 2B**). Earlier work reported that Rhi depletion leads to loss of UAP56 at piRNA cluster loci (Zhang et al. 2012). Thus, we probed a potential role of CG13741 and Nxf3 in the recruitment of UAP56 (**Fig. 2C**). As expected, UAP56 recruitment depended on all components of the RDC complex, and it also depended on CG13741. In contrast, germline knockdowns of *nxf3* had no effect on the localisation of UAP56, suggesting that CG13741 acts upstream of both Nxf3 and UAP56 (**Fig. 2C**).

Of note, germline knockdown of *moon* led to pronounced accumulation of CG13741, Nxf3 and UAP56 in nuclear foci that partially co-stained for Rhi (**Fig. 2A-C**). Presently, the nature of such loci is unclear.

Depletion of piRNA dual-strand cluster factors did not eliminate nuclear Piwi signal (**Fig. 2B**) nor did depletion of Piwi from ovarian germ cells impact on factors involved in dual-strand piRNA cluster transcription, as reported (Akkouche et al. 2017). Conversely, knockdown of *piwi* did not alter the localisation of CG13741, UAP56 or Nxf3, suggesting that Piwi is not required, at least in the short term, for formation of the signals that recruit these proteins to cluster loci (**Fig. 2A-C**). Lastly, Vas localization was not affected by loss of any of the factors studied, consistent with its role in assembling nuage and a potential function downstream of piRNA cluster transcription and export (**Fig. 2C**).

The role of UAP56 in Nxf3 and CG13741 recruitment was also tested using *uap56*^*sz15’/28*^ mutants. Only UAP56 localisation at clusters is affected in these mutants. Nxf3 and CG13741 levels were strongly reduced in the mutants compared to controls, while weak signals were still detected at discrete nuclear foci (**Fig. S2B**). Furthermore, CG13741 and Rhi colocalization was maintained in *uap56*^*sz15’/28*^ mutants (**Fig. S2B, left**).

Considered together, our data indicate that CG13741 and Nxf3 recruitment is dependent on the presence of the RDC complex. CG13741 is essential for the correct localization of both Nxf3 and UAP56 to dual-strand piRNA clusters (**Fig. 2D,E**), whereas Nxf3 in turn could promote translocation of some CG13741 protein outside of the nucleus. Given the involvement of CG13741 in the biology of dual-strand piRNA clusters, we have, by mutual agreement, named CG13741 Bootlegger (ElMaghraby et al. 2019).

### Bootlegger and Nxf3 are required for piRNA production from dual-strand clusters

To explore the genome-wide impact of *bootlegger* and *nxf3* mutations, we performed RNA-seq from ovaries of mutant flies and compared these to control flies. The expression of protein-coding genes was only mildly affected in *bootlegger*^*mut*^ (r^2^ = 0.96) or *nxf3*^*mut*^ (r^2^ = 0.99), while loss of Bootlegger or Nxf3 led to a strong de-repression of germline (*burdock, HeT-A* and *TAHRE*) and intermediate (*blood*) transposons (**Fig. 3A**). Somatic transposons were not changed in *bootlegger* and *nxf3* mutants, as expected based upon their expression patterns (**Fig. 3A**). 14 out of 60 germline transposon families (above the expression threshold) showed more than 4-fold upregulation in *bootlegger* mutants (**Fig. 3A, top**). In *nxf3* mutants, 10 out 60 germline transposons were de-repressed more than 4-fold (**Fig. 3A, bottom**), and these defects in transposon silencing were comparable to those observed in *rhi and moon* mutants (**Fig. S2C, left and middle**). Notably, *bootlegger* and *nxf3* mutants had very similar impacts overall, considering effects on genes and transposons (**Fig. S2C, right**).

**Figure 3.**
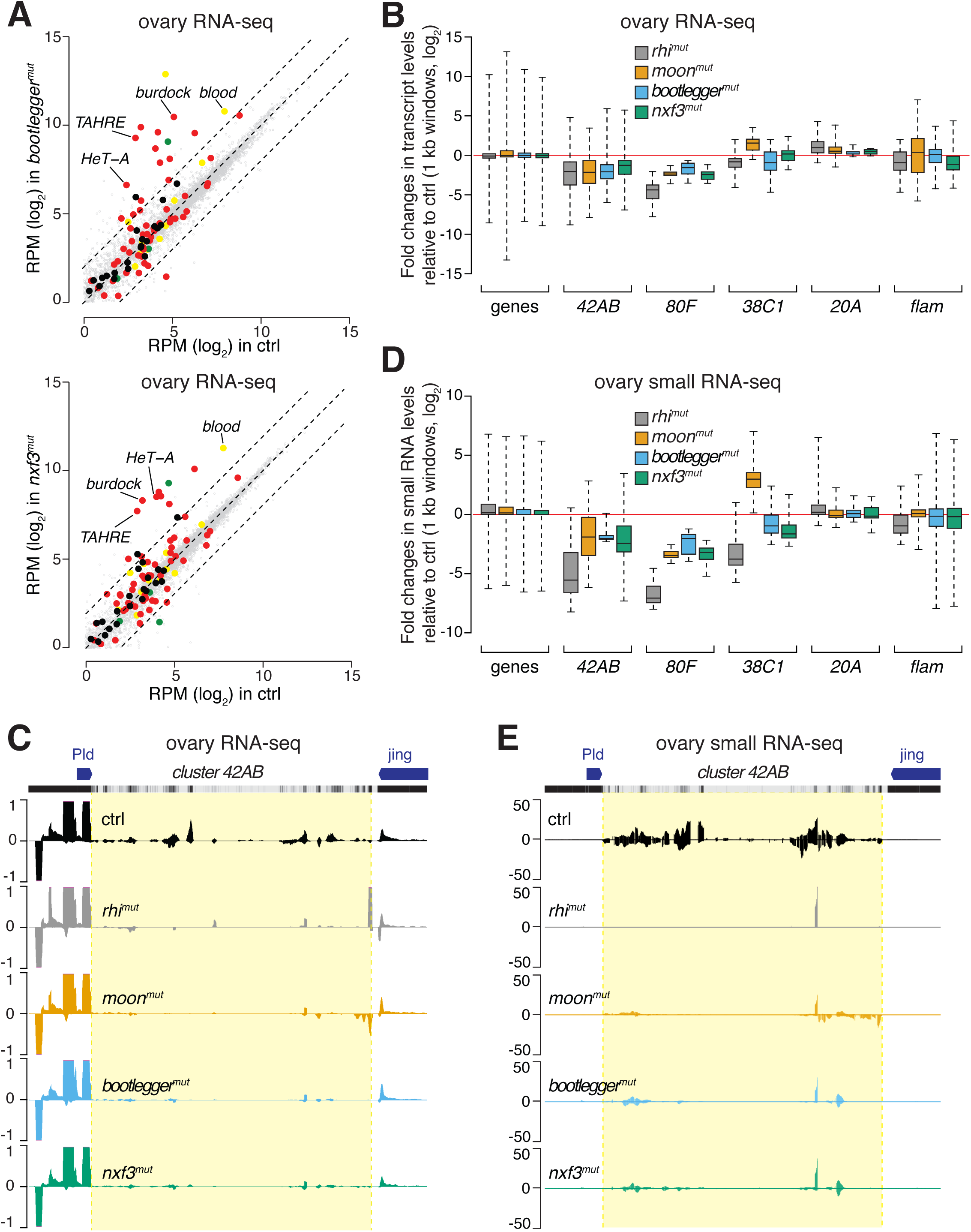
Bootlegger and Nxf3 are essential for piRNA production. **(A)** Scatter plots showing expression levels (reads per million sequenced reads, RPM) of genes (in grey) and transposons (red, germline-specific; yellow, intermediate; green, soma-specific; black, other transposons) from total RNA from ovaries of the indicated genotypes (top, *bootlegger*^*mut*^; bottom, *nxf3*^*mut*^; n = 3) compared to control flies. Dashed lines indicate 4-fold expression changes. **(B)** Box plots showing changes in RNA levels between the indicated genotypes and control in ovarian RNA-seq for the indicated categories (1 kb bins for piRNA cluster transcripts *42AB, 80F, 38C1, 20A, flam*). Grey, *rhi*^*mut*^; orange, *moon*^*mut*^; blue, *bootlegger*^*mut*^; green, *nxf3*^*mut*^. **(C)** UCSC genome browser shot displaying profiles of RNA-seq levels of reads uniquely mapping to cluster *42AB* and flanking euchromatic regions in the indicated genotypes. Black, control; grey, *rhi*^*mut*^; orange, *moon*^*mut*^; blue, *bootlegger*^*mut*^; green, *nxf3*^*mut*^. The mappability of 50 bp reads is shown above. **(D)** As in (B) but shown are box plots of small RNA-seq for the indicated categories (1 kb bins for piRNA cluster transcripts *42AB, 80F, 38C1, 20A, flam*). Grey, *rhi*^*mut*^; orange, *moon*^*mut*^; blue, *bootlegger*^*mut*^; green, *nxf3*^*mut*^. **(E)** As in (C) but shown are piRNAs (23 to 29-nt size) uniquely mapping to cluster *42AB* and flanking euchromatic regions in the indicated genotypes. Black, control; grey, *rhi*^*mut*^; orange, *moon*^*mut*^; blue, *bootlegger*^*mut*^; green, *nxf3*^*mut*^. The mappability of 25 bp reads is shown above.

Previous reports showed that loss of the RDC complex results in down-regulation of piRNA precursors derived from dual-strand clusters (Mohn et al. 2014; Zhang et al. 2014). To evaluate the potential effects of *bootlegger* and *nxf3* mutations on piRNA cluster levels, we extracted reads uniquely mapping to piRNA source loci and binned them into 1 kb windows (see methods for details). Loss of either Bootlegger or Nxf3 had a pronounced impact on the levels of piRNA precursors derived from dual-strand clusters, though the effects were milder than those observed in *rhi* mutants (**Fig. 3B**). Interestingly, the levels of piRNA precursors from cluster *42AB* were reduced more severely in *bootlegger* mutant ovaries then they were in *nxf3* knockouts (**Fig. 3B,C**). These data are in line with Bootlegger acting upstream of Nxf3 and suggest a function of Bootlegger in cluster transcription or transcript stability in addition to its role in recruitment of Nxf3. In contrast, the reduction of RNA from piRNA cluster *80F* was comparable between *bootlegger* and *nxf3* mutants (**Fig. S2D).** Of note, the levels of transcripts derived from the uni-strand clusters, *flam* and *20A*, remained unchanged (**Fig. 3B; Fig. S2D**).

We sequenced small RNA libraries prepared from total RNA from ovaries of *bootlegger* and *nxf3* mutant flies and observed a global reduction in repeat-derived piRNAs in both mutants (**Fig. S3A**). We specifically extracted reads mapping to piRNA source loci using the same binning strategy described above. Levels of piRNAs mapping to the major dual-strand clusters were severely reduced in *bootlegger* and *nxf3* mutants (**Fig. 3D,E; Fig. S3B**). Compared to wild-type controls, we observed a strong reduction of sense and antisense piRNAs from cluster *42AB* upon loss of either Bootlegger or Nxf3, similar to *moon* mutants but weaker than *rhi* mutants (**Fig. 3D,E**). In contrast, piRNAs mapping to cluster *20A* and *flam* remained unchanged, as were gene-derived piRNAs (**Fig. 3D; Fig. S3B**). Considered together, our data are consistent with a model in which Nxf3 and Bootlegger are specifically required for the production of piRNAs from dual-strand cluster transcripts.

### Nxf3 promotes export of piRNA cluster transcripts to cytoplasmic processing sites

Unlike RDC components and Moon, Bootlegger and Nxf3 could be detected outside nuclei in addition to their presence in nuclear foci (**Fig. 1D,E**). We confirmed their perinuclear localisation by co-staining with Vas, which marks nuage in nurse cell nuclei (Lim and Kai 2007). While most Nxf3 foci were detected in nuclei, some signal overlapped with Vas staining (**Fig. 4A; Fig. S4A**). The same pattern was observed for Bootlegger, whereas Rhi was present only in nuclear foci (**Fig. 4A; Fig. S4A**), as previously reported (Klattenhoff et al. 2009; Mohn et al. 2014; Zhang et al. 2014). These results suggest that Bootlegger and Nxf3 transit between the nucleus and cytoplasm, where they are present in nuage, the sites of piRNA biogenesis in germ cells.

**Figure 4.**
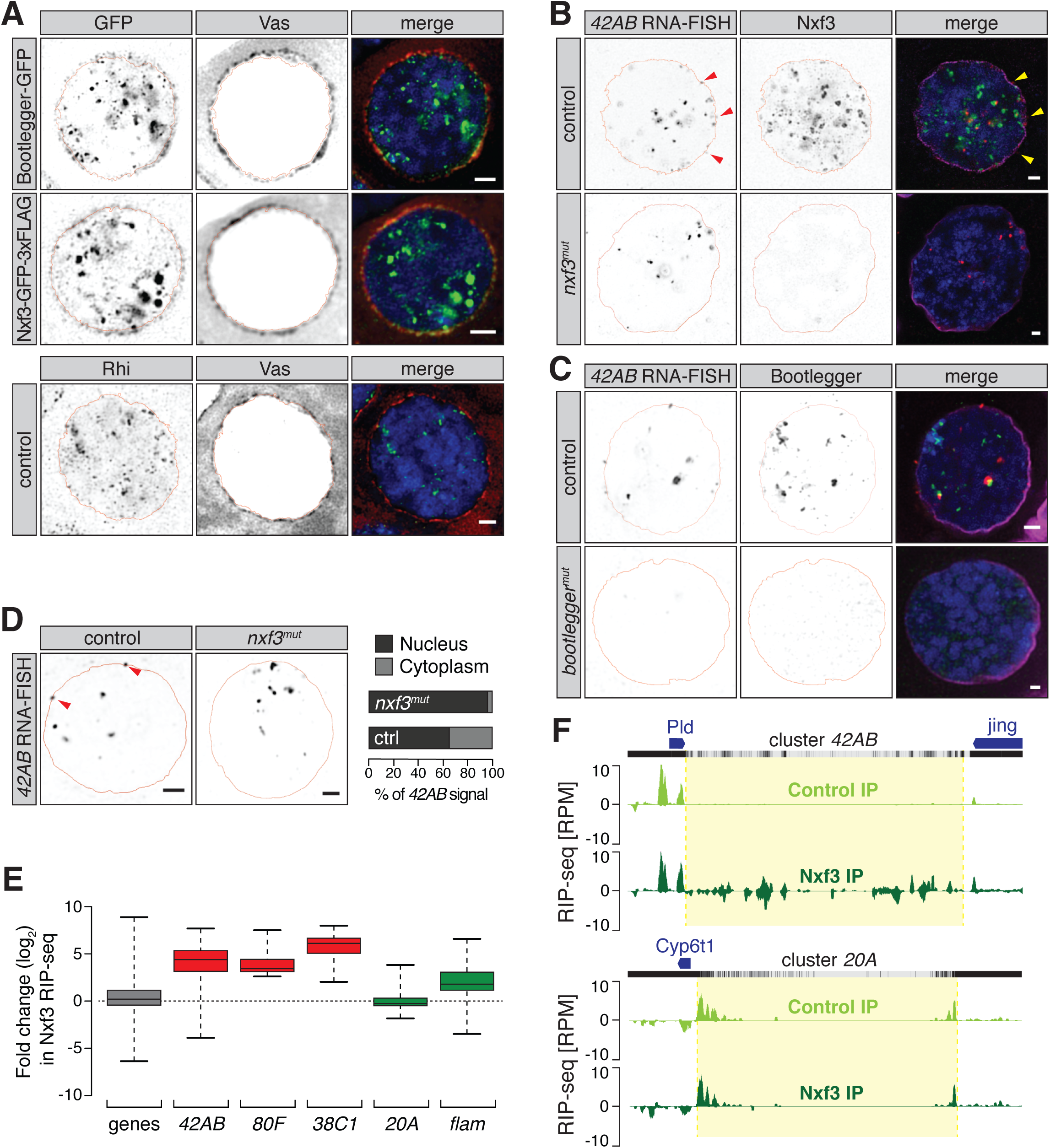
Nxf3 promotes dual-strand piRNA cluster export. **(A)** Expression and localization of Bootlegger-GFP (top), Nxf3-GFP-3xFLAG (middle), and Rhi in nurse cell nuclei is shown by immunofluorescence and co-stained with Vas. Green, GFP/Rhi; red, Vas; blue, DNA. Scale bar, 2 µm. **(B)** Expression and localization of Nxf3 protein and cluster *42AB* RNA are shown by immunofluorescence and RNA-FISH in control and *nxf3*^*mut*^ nurse cell nuclei. Green, Nxf3; red, cluster *42AB* RNA; magenta, Lamin, blue, DNA. Scale bar, 2 µm. **(C)** As in (B) but showing expression and localization of Bootlegger protein and cluster *42AB* RNA in control and *bootlegger*^*mut*^ nurse cell nuclei. Green, Bootlegger; red, cluster *42AB* RNA; magenta, Lamin, blue, DNA. Scale bar, 2 µm. **(D)** Expression and localization of piRNA cluster *42AB* transcripts are shown by RNA-FISH in control and *nxf3*^*mut*^ nurse cell nuclei (left). Scale bar, 2 µm. The number of nuclear and cytoplasmic foci was quantified for control (n = 92) and *nxf3* mutant (n = 97) nurse cell nuclei (right). **(E)** Box plots showing changes in Nxf3 RIP-seq levels compared to control for the indicated categories (1 kb bins for piRNA cluster transcripts *42AB, 80F, 38C1, 20A, flam*). Grey, protein-coding genes; red, dual-strand clusters; green, uni-strand clusters. **(F)** UCSC genome browser shot displaying profiles of Nxf3 RIP-seq levels of reads uniquely mapping to cluster *42AB* (top) and cluster *20A* (bottom) in the indicated conditions. Light green, control IP; dark green, Nxf3 IP. Shown are reads per million (RPM). The mappability of 50 bp reads is shown above.

Given that Nxf3 is part of the mRNA nuclear export factor family, the localization pattern of Nxf3 protein in nuclei and nuage, and the effects of *nxf3* loss on cluster transcripts and mature piRNAs, suggested a role for Nxf3 in the export of piRNA precursors. To test this hypothesis, we performed RNA-FISH for cluster *42AB* transcripts in wild-type control, *nxf3*^*mut*^, and *bootlegger*^*mut*^ ovaries. No signal was detected in nurse cells of *bootlegger* mutants, whereas *nxf3*^*mut*^ ovaries retained only staining of nuclear foci (**Fig. 4B,C; Fig. S4B**). In control ovaries, the majority of the signal detected corresponded to nuclear foci, however, we also detected cytoplasmic foci (typically 1-2 foci per stack), corresponding to exported precursors (**Fig. 4B,C; Fig. 4D, left**). While *nxf3* mutants did not affect the nuclear foci, we observed a substantial reduction of cluster *42AB* transcript signals outside of the nuclei (**Fig. 4D**). In control flies, 35% of cluster *42AB* RNA signal was cytoplasmic, compared to only 4% of signal in the cytoplasm in *nxf3* mutants, highlighting a critical function of Nxf3 in piRNA precursor export (**Fig. 4D, right)**.

Given the implication of Nxf3 function in dual-strand cluster export, we tested whether there was a direct interaction between Nxf3 and cluster transcripts. We immunoprecipitated Nxf3 protein from control and *nxf3*^*mut*^ ovaries and analyzed the associated RNA by RIP-seq (RNA immunoprecipitation sequencing). We detected a strong enrichment for RNAs derived from dual-strand clusters in Nxf3 RIP, whereas transcripts of protein-coding genes were not enriched in comparison to the control (**Fig. 4E**). Transcripts from cluster *42AB* and cluster *80F* were enriched ∼20-fold, while the levels of the uni-strand cluster *20A* were comparable between control and mutant ovaries (**Fig. 4E,F; Fig. S4C**). These data indicate that Nxf3 binds predominantly to piRNA precursor transcripts derived from dual-strand clusters in nurse cells.

### Nxf3-mediated piRNA precursor export requires Nxt1, Bootlegger and Crm1

Eukaryotic cells employ several strategies to ensure proper mRNA export (Bjork and Wieslander 2017). Canonical mRNA export by Nxf1 depends on heterodimer formation with its cofactor Nxt1 and this interaction is mediated through their NTF2-like domains (Herold et al. 2001; Kohler and Hurt 2007; Stewart 2010). Nxf3 is also predicted to contain a NTF2-like domain (**Fig. 1B**), thus we probed for a potential interaction between Nxf3 and Nxt1. In wild-type ovaries, Nxf3 and Nxt1 colocalized to distinct foci of nurse cell nuclei (**Fig. 5A**), while Nxt1 showed additional, dispersed signal throughout the nucleus. In *nxf3*^*mut*^ ovaries, all nuclear Nxt1 foci were lost while the disperse signal remained unchanged (**Fig. 5A**). Next, we analyzed the localization of Nxf3 upon *nxt1* germline knockdown. Depletion of Nxt1 leads to severe morphological defects of the ovaries, likely due to its function in general mRNA export (**Fig. 5B**). To identify the germ cells in Nxt1 depleted ovaries, we used Vas staining, and found that Nxf3 was still detected in germ cell nuclei (**Fig. 5B**). Thus, our data are consistent with a model in which Nxf3 and Nxt1 likely form a complex in nurse cell nuclei, with Nxf3 recruiting Nxt1 to piRNA cluster transcripts. To test whether Nxf3 and Nxt1 interact, we expressed both proteins in S2 cells. Co-immunoprecipitation and western blot analysis showed an association between Nxf3 and Nxt1 (**Fig. 5C, top**). Since Bootlegger recruitment to piRNA clusters depends on RDC (**Fig. 2A**), and Nxf3 fails to associate with RDC foci in the absence of Bootlegger (**Fig. 2B**), we reasoned that Bootlegger could be responsible for bringing Nxf3 to piRNA precursors via an interaction between these proteins. Indeed, Nxf3 and Bootlegger co-immunoprecipitated in S2 cells and reciprocal co-immunoprecipitation from ovaries confirmed that Bootlegger and Nxf3 interact *in vivo* (**Fig. 5C, bottom; Fig. S4D**), leading us to conclude that Bootlegger recruits Nxf3, which subsequently brings Nxt1, to dual-strand clusters, ultimately linking RDC-dependent cluster transcription with Nxf3-dependent transcript export.

**Figure 5.**
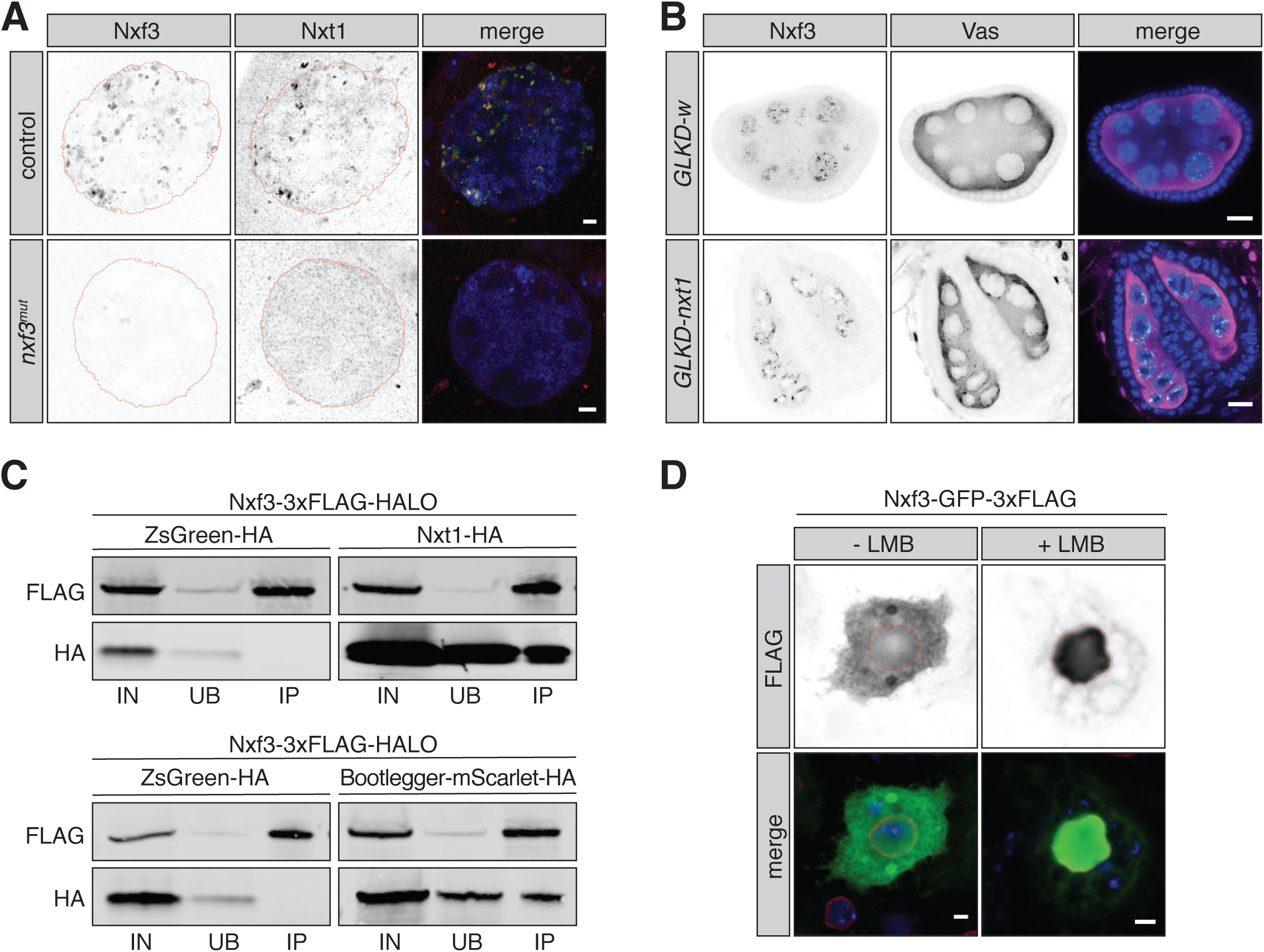
Nxf3 interacts with Nxt1 at piRNA cluster loci and piRNA precursor export requires Crm1. **(A)** Expression and localization of Nxf3 and Nxt1 in control and *nxf3*^*mut*^ nurse cell nuclei is shown by immunofluorescence. Green, Nxf3; red, Nxt1; blue, DNA. Scale bar, 2 µm. Expression and localization of Nxf3 and Vas in egg chambers upon germline-specific knockdown (GLKD) of *nxt1* or *w* (control) are shown by immunofluorescence. Green, Nxf3; magenta, Vas; blue, DNA. Scale bar, 10 µm. **(C)** Western blot analyses of FLAG-tag pulldown from lysates of S2 cells transfected with the indicated expression constructs. IN, input (10%); UB, unbound (2.5%); IP, immunoprecipitate (50%). **(D)** Expression and localization of the Nxf3-GFP-3xFLAG construct in S2 cells treated with leptomycin B (LMB) or control treatment are shown by immunofluorescence. Green, FLAG; red, Lamin; blue, DNA. Scale bar, 2 µm.

Nxf1 is responsible for canonical mRNA export, however, the absence of key mRNA processing such as splicing and polyadenylation led us to investigate alternative export mechanisms for piRNA precursor transcripts. The nuclear exportin Crm1 is largely responsible for the export of proteins, but has also been reported to assist in nuclear export of RNAs via some NXF family proteins, including human Nxf3 (Yang et al. 2001; Kohler and Hurt 2007). Moreover, both mouse and *Xenopus* Nxt1 were shown to directly bind Ran-GTP and to activate Crm1-dependent nuclear export (Ossareh-Nazari et al. 2000; Black et al. 2001), leading us to postulate that Crm1 might facilitate Nxf3-Nxt1-dependent export of piRNA precursors from *Drosophila* nurse cell nuclei. To test this hypothesis, we exploited a small molecule drug, Leptomycin B (LMB), which specifically blocks Crm1-dependent export (Kudo et al. 1999). We transfected S2 cells with a Nxf3-GFP-3xFLAG expression construct and treated the cells with LMB for 12 hours. In the absence of LMB, the Nxf3 fusion protein was uniformly distributed in nuclei and the cytoplasm (**Fig. 5D**). In contrast, treatment with LMB resulted in accumulation of Nxf3-GFP-3xFLAG in nuclei (**Fig. 5D**). These results suggest that Nxf3 contains an intrinsic nuclear localization signal but can be shuttled to the cytoplasm. Considered together, our data indicate that Nxf3-mediated export of piRNA precursors depends on Crm1.

## Discussion

The transcription and export of piRNA precursor transcripts requires a highly specialized machinery that must assemble correctly at dual-strand cluster loci, initiate non-canonical transcription, and license and transport these transcripts (which lack features of processed and export-competent mRNAs) to the cytoplasm where they are processed into mature piRNAs. How each step is achieved and how the elements involved interact is yet to be fully understood.

Here we show that export of piRNA precursors from dual-strand clusters in nurse cells depends on a specific mechanism that requires Bootlegger, a protein without known domains, and the nuclear export factor Nxf3 (**Fig. 6**). We find that Bootlegger is important for either the synthesis or stability of transcripts from dual-strand piRNA clusters and is required for Nxf3 recruitment to dual-strand piRNA cluster loci. Analysis of RNA-seq, small RNA-seq, and RIP-seq for Nxf3, combined with immunofluorescence and RNA-FISH analyses provide evidence that Nxf3 binds to and transports piRNA precursor transcripts from their sites of transcription (piRNA clusters) to the sites where piRNA processing takes place (nuage). We further find that Nxf3-mediated export depends on Crm1. These findings are all in agreement with those of ElMaghraby and colleagues who also identified Nxf3 and Bootlegger as critical facilitators of piRNA precursor export (ElMaghraby et al. 2019).

**Figure 6.**
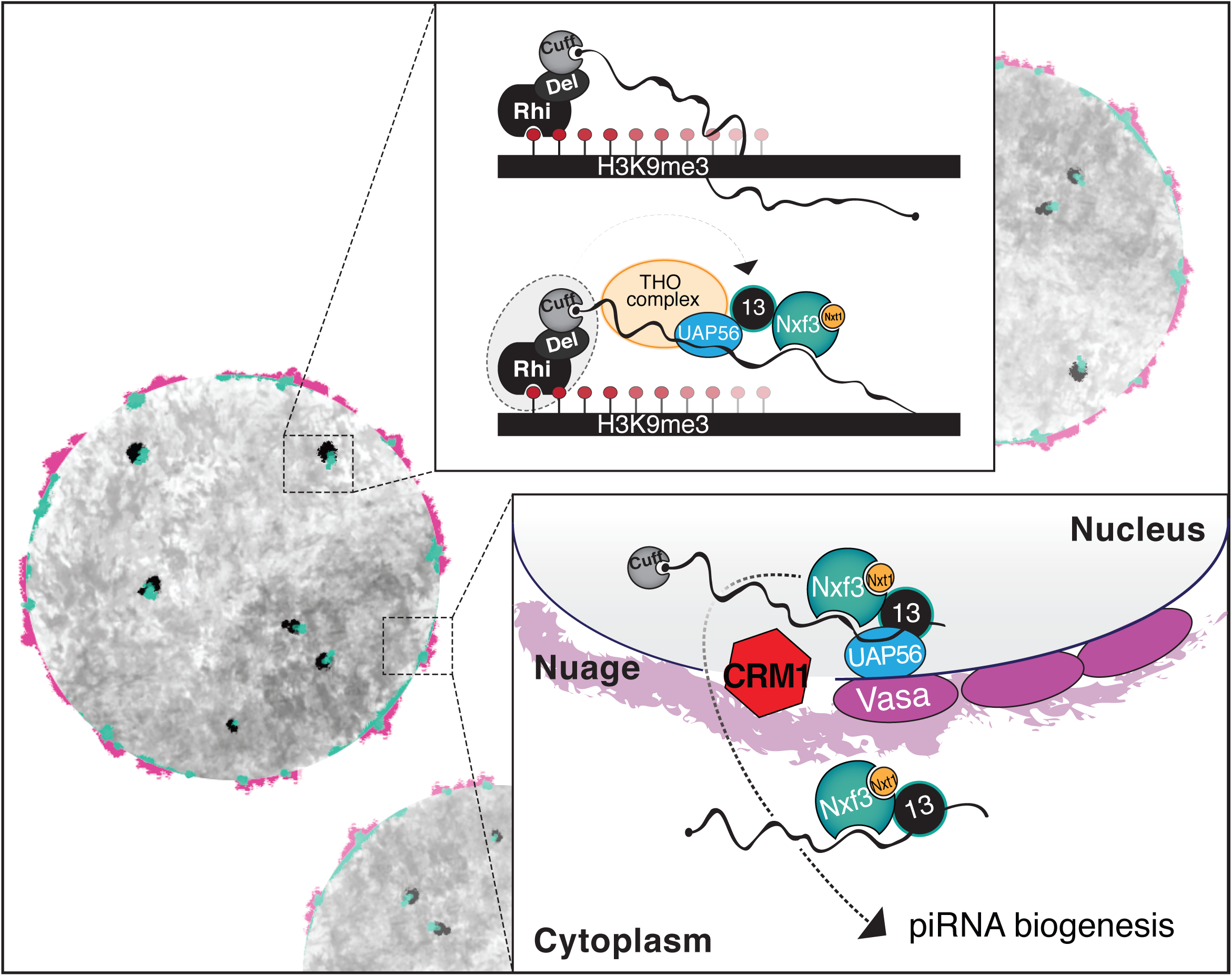
Model of piRNA cluster transcript generation and export. In germ cells nuclei of the *Drosophila* ovary, the RDC complex licenses dual-strand piRNA cluster transcription. Bootlegger is recruited to cluster loci via RDC components. Bootlegger instruments the recruitment of UAP56 followed by the THO complex and Nxf3 [top inset]. Nxf3 associates with piRNA precursors and Nxt1. Subsequently, Bootlegger, Nxt1, and Nxf3 in complex with piRNA precursors transit to nuage in a Crm1-dependant process. Following their export, dual-strand piRNA cluster transcripts are funnelled into the piRNA biogenesis machinery [bottom inset].

How Bootlegger is recruited to piRNA cluster loci remains to be determined. One plausible mechanism is via a direct protein-protein interaction with one or more components of the RDC complex, which specifies piRNA clusters upon recognition of H3K9me3 marks by Rhi. In support of this hypothesis, an interaction between Bootlegger and Del was observed by a yeast-2-hybrid screen (ElMaghraby et al. 2019), however, whether this complex forms in ovaries requires further examination. Recruitment of Nxf3, in turn, requires Bootlegger, likely also through direct protein-protein interaction. It will be important to uncover the precise contacts that drive this recruitment and to probe if RNA binding by Nxf3 (and/or Bootlegger) might contribute to complex formation. Nxf3 likely binds cluster RNAs through its cargo-binding domain, which is composed of an RNA-binding domain and LRRs. But how specific binding of piRNA precursors is achieved remains a mystery. In fact, how exclusivity for the three NXF proteins with annotated functions (Nxf1-3) is achieved is a critical question for the future.

Eukaryotic cells employ various different RNA export machineries, depending on the class of cargo RNA. Canonical mRNAs, which carry 5’ caps, have undergone splicing, and carry 3’ poly(A) tails are exported via the Nxf1-Nxt1 pathway. Another extreme example are pre-miRNAs, which are specifically recognized via their secondary structure and exported by Exportin-5 (Yi et al. 2003). A subset of RNAs are exported via a Crm1 (Chromosomal maintenance 1)-dependant mechanism. Although Crm1 is typically involved in protein export, it can function in RNA export via adapter proteins.

For example, Crm1 binds to the Cap-binding complex and the adapter protein PHAX to export ∼200-nt sized small nuclear RNAs (Ohno et al. 2000). Here we show that Nxf3-mediated export of piRNA precursor transcripts also requires Crm1. The role of Nxt1 in piRNA precursor export is not yet clear. It is plausible that Nxt1 is required for export by Nxf3, or it might function in co-transcriptional recruitment to cargo RNAs. In this regard, it is striking that Nxt1, and also UAP56 and THO complex components are concentrated at piRNA clusters (Zhang et al. 2012; Zhang et al. 2018), even though these proteins are reportedly essential for general mRNA export.

The work presented here, and other emerging realizations (Batki et al. 2019; ElMaghraby et al. 2019; Fabry et al. 2019; Murano et al. 2019; Zhao et al. 2019) begin to paint a picture in which the evolutionary pressures exerted by the need to control mobile genetic elements have resulted in exaptation and dedication of nuclear export factor family members to the piRNA pathway. This specialization has taken different forms. The most easily understood is the adaptation of Nxf3 to export a specific class of non-canonically transcribed RNAs (ElMaghraby et al. 2019), whereas the precise mechanism by which Nxf2 acts as a co-transcriptional silencing factor remains more mysterious (Batki et al. 2019; Fabry et al. 2019; Murano et al. 2019; Zhao et al. 2019). Ultimately, an understanding of how these proteins have assumed new roles through evolution will require a detailed understanding of how they are recruited to particular transcripts and how they mediate their downstream effects.

## Acknowledgements

We thank Martin H. Fabry for help with computational analyses. We thank the CRUK Cambridge Institute Bioinformatics, Genomics, Microscopy, Research Instrumentation and Cell Services, and Proteomics Core Facilities for technical support. We thank the University of Cambridge Department of Genetics Fly Facility for microinjection services and fly stock generation. We thank the Vienna *Drosophila* Resource Center and the Bloomington Stock Center for fly stocks. We thank Julius Brennecke for anti-Piwi, anti-Aub and anti-Ago3 antibodies, William Theurkauf for anti-Rhi antibody and fly strains, Elisa Izaurralde for anti-Nxt1 antibody, and Paul Lasko for anti-UAP56 antibody. Early work on this project was funded by HHMI and by a kind gift from Kathryn W. Davis at Cold Spring Harbor Laboratory. Research in the Hannon laboratory is supported by Cancer Research UK and by a Wellcome Trust Investigator award 110161/Z/15/Z. M.M. is supported by a Boehringer Ingelheim Fonds PhD fellowship.

## Author contributions

E.K. performed all experiments with help from M.M., E.L.E., U.-S.D. and B.C.. J.B.P. generated the CG13741 antibody and a CG13741 construct. E.K., G.J.H. and B.C. designed the experiments, analyzed and interpreted the data, and wrote the manuscript with input from M.M. and E.L.E.

## Supplementary Figure legends

**Figure S1.**
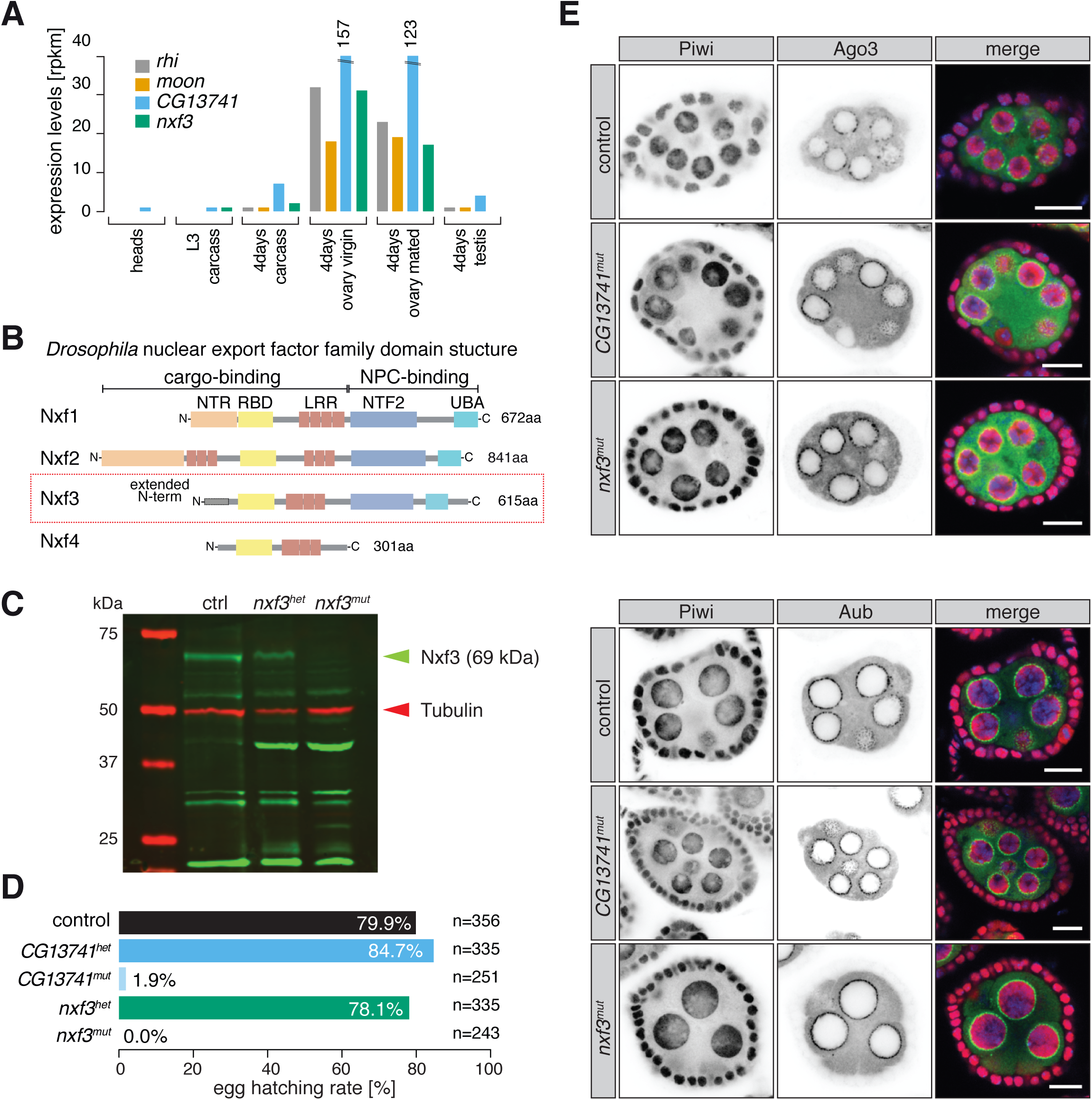
**(A)** Bar graph showing expression levels of *rhi, moon, CG13741, nxf3* in various *Drosophila* developmental stages and tissues. **(B)** Schematic representation of Nxf1, Nxf2, Nxf3, Nxf4 domain organizations. NTR, N-terminal region; RBD, RNA binding domain; LRR, Leucine reach repeat; NTF2, NTF2-like domain; UBA, Ubiquitin-associated domain; NPC, nuclear pore complex. **(C)** Nxf3 expression levels in *nxf3* mutant, *nxf3* heterozygote and control ovaries were analyzed by western blot. **(D)** Percentage of eggs hatching from female flies from the indicated genotypes. **(E)** Localization of Piwi, Aub and Ago3 in control, *nxf3*^*mut*^ and *CG13741*^*mut*^ flies in an egg chamber are shown by immunofluorescence. Green, Ago3/Aub; red, Piwi; blue, DNA. Scale bar, 10 µm.

**Figure S2.**
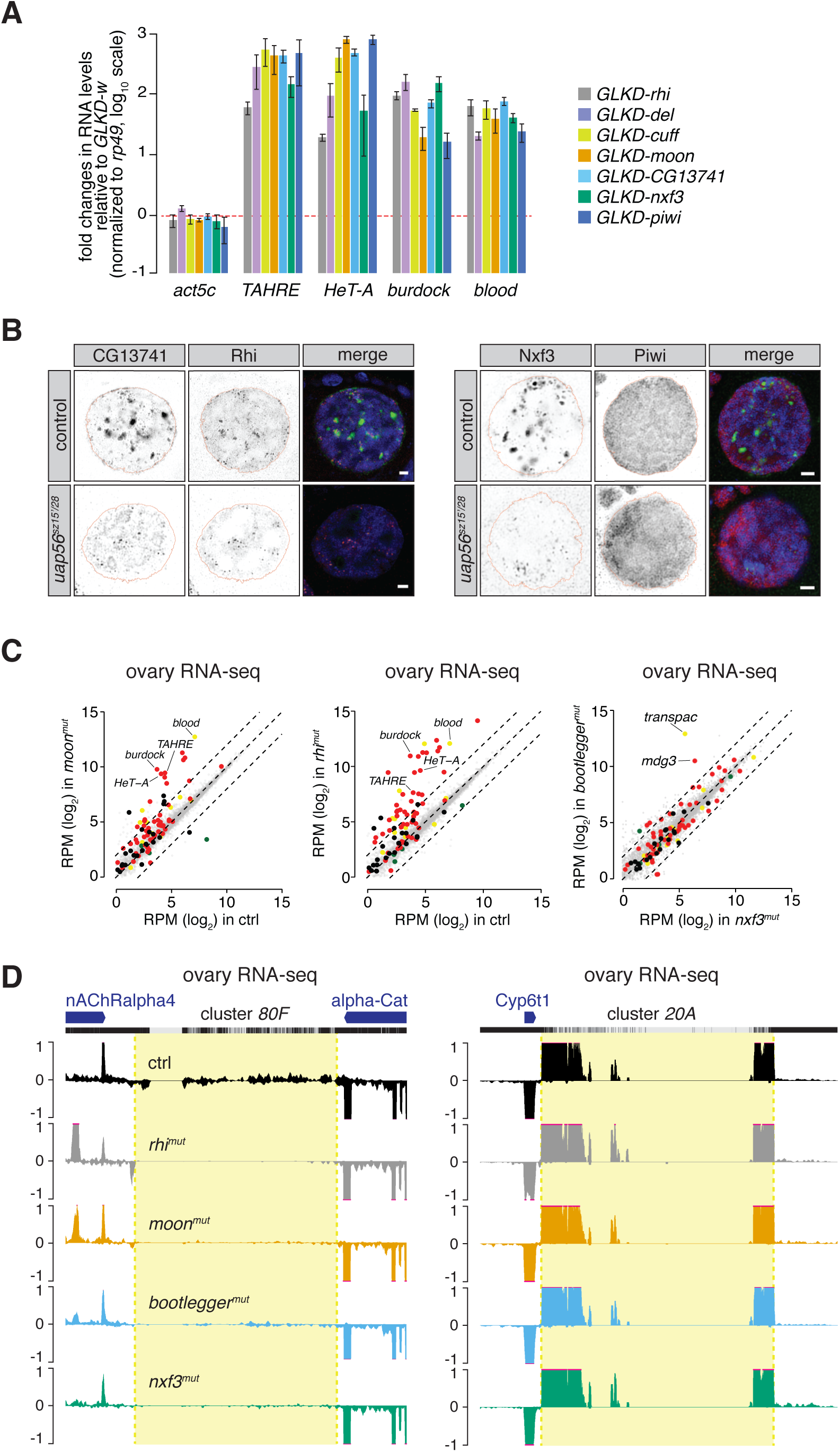
**(A)** Transposon levels were measured by qPCR in germline knockdowns of *rhi, del, cuff, moon, piwi, nxf3* and *CG13741* and compared to depletion of *w*. Results were normalized to *rp49*. Error bars indicate standard deviation (n = 3). **(B)** Expression and localization of Rhi, CG13741, Nxf3 and Piwi in nurse cell nuclei of control and *uap56*^*sz15*^*/uap56*^*28*^ mutant flies is shown by immunofluorescence. Green, CG13741/Nxf3; red, Rhi/Piwi; blue, DNA. Scale bar, 2 µm. **(C)** Scatter plots showing expression levels (reads per million sequenced reads, RPM) of genes (in grey) and transposons (red, germline-specific; yellow, intermediate; green, soma-specific; black, other transposons) from total RNA from ovaries of the indicated genotypes (left, *moon*^*mut*^ compared to control; middle, *rhi*^*mut*^ compared to control; right, *bootlegger*^*mut*^ compared to *nxf3*^*mut*^; n = 3). Dashed lines indicate 4-fold expression changes. **(D)** UCSC genome browser tracks showing the coverage of reads uniquely mapping to cluster *80F* (left) and *20A* (right) for the indicated genotypes. The mappability tracks calculated for 50 bp is shown above. Black, control; grey, *rhi*^*mut*^; orange, *moon*^*mut*^; blue, *bootlegger*^*mut*^; green, *nxf3*^*mut*^.

**Figures S3.**
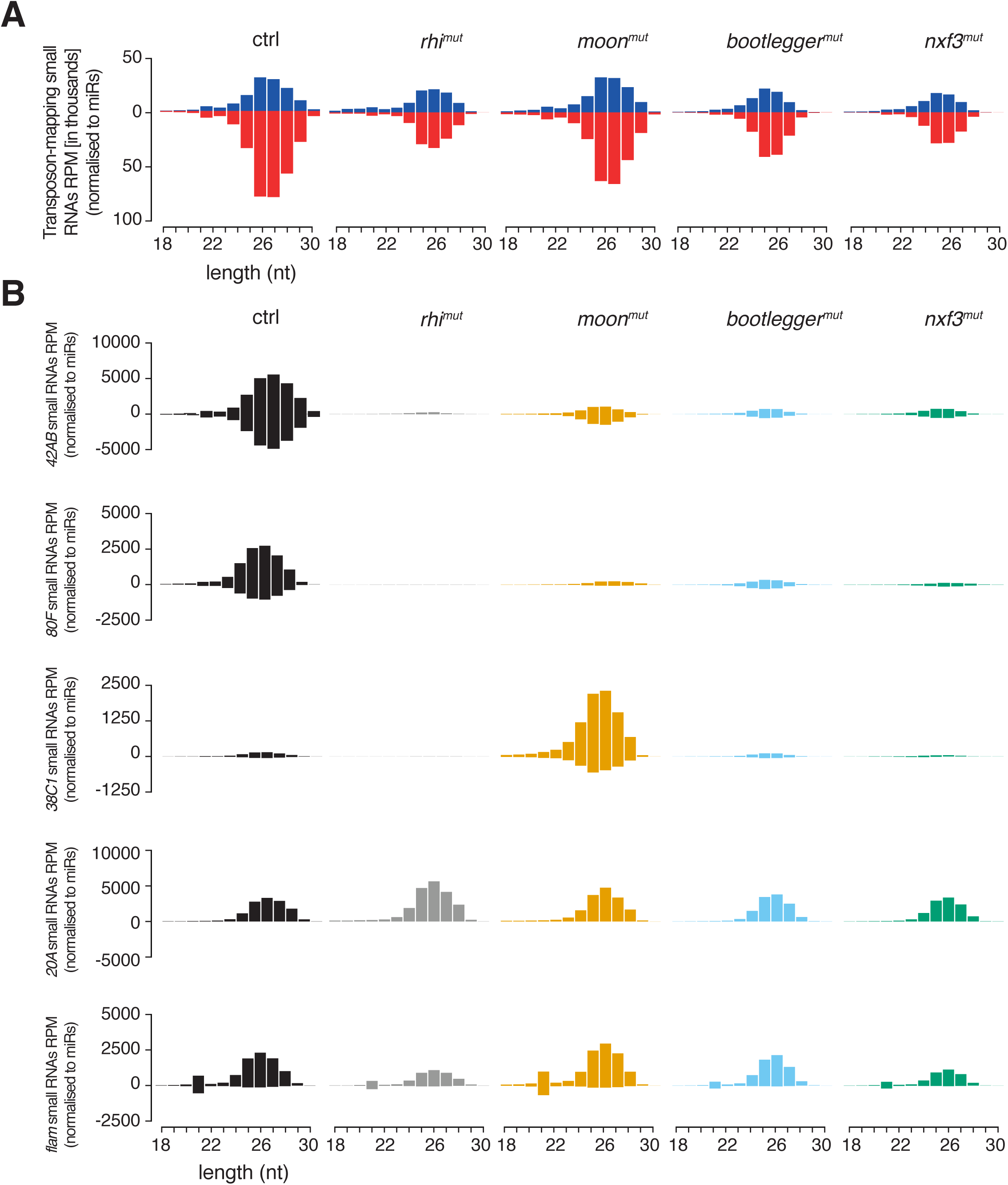
**(A)** Bar graphs showing the size distribution of transposon-mapping small RNAs from ovaries for the indicated genotypes. **(B)** Bar graphs showing the size distribution of cluster mapping small RNAs from ovaries for the indicated genotypes and clusters.

**Figures S4.**
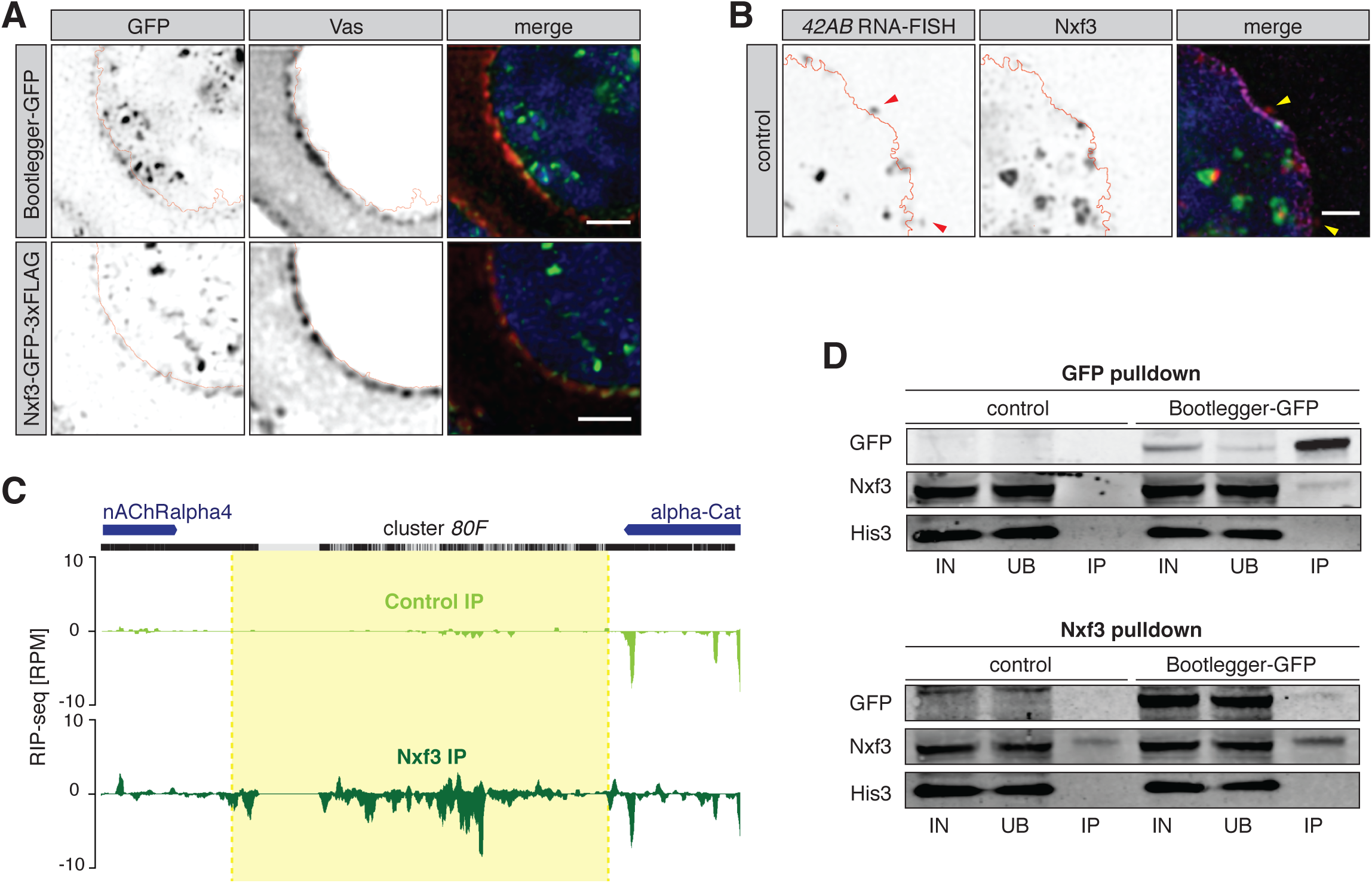
**(A)** Expression and localization of Bootlegger-GFP (top), Nxf3-GFP-3xFLAG (middle), and Rhi in zoom-in of nurse cell nuclei is shown by immunofluorescence and co-stained with Vas. Green, GFP; red, Vas; blue, DNA. Scale bar, 2 µm **(B)** Expression and localization of Nxf3 protein and cluster *42AB* RNA in zoom-ins of nurse cell nuclei are shown by immunofluorescence and RNA-FISH. Green, Nxf3; red, cluster *42AB* RNA; magenta, Lamin; blue, DNA. Scale bar, 2 µm. **(C)** UCSC genome browser shot displaying profiles of Nxf3 RIP-seq levels of reads uniquely mapping to cluster *80F* in the indicated conditions. Light green, control IP; dark green, Nxf3 IP. Shown are reads per million (RPM). The mappability of 50 bp reads is shown above. **(D)** Western blot analyses of GFP or Nxf3 pulldown from ovarian lysates of Bootlegger-GFP or control flies. IN, input; UB, unbound; IP, immunoprecipitate.

## Supplementary Tables

**Table S1. Fly strains.**

**Table S2. qPCR primers and gRNAs.**

**Table S3. Amino acidic sequence of extended Nxf3.**

## Materials and Methods

### Fly stocks and handling

All flies were kept at 25 °C on standard cornmeal or propionic food. CG13741-GFP, extended Nxf3-GFP-3xFLAG, annotated Nxf3-GFP-3xFLAG, *CG13741* mutant alleles (*CG13741*^*Δ1*^ and *CG13741*^*Δ2*^) and the *nxf3* mutant allele (*nxf3*^*mut*^) were generated for this study (see below). Control *w*^*1118*^ flies were a gift from the University of Cambridge Department of Genetics Fly Facility. For germline-specific knockdowns we used a stock containing a UAS::Dcr2 transgene and a nos::GAL4 driver (Czech et al. 2013) and shRNA and dsRNA lines from the Bloomington Drosophila Stock Center and Vienna Drosophila Resource Center. Fertility of mutant females was scored by crossing ten freshly hatched females to five *w*^*1118*^ males and counting the number of eggs laid in 12 hr periods and pupae that developed after 7 days. Details on all fly stocks used in this study are listed in **Table S1**.

### Generation of mutant and transgenic fly strains

Frameshift mutant alleles of *CG13741 and nxf3* were generated by injecting pCFD4 (a gift from Simon Bullock; Addgene plasmid # 49411 ; http://n2t.net/addgene:49411; RRID:Addgene_49411; (Port et al. 2014)) containing two gRNAs against *CG13741* and *nxf3* into embryos expressing vas::Cas9 (Bloomington stock 51323). CG13741-GFP was cloned in an in-house generated transgenesis vector for phiC31-mediated integration and expressed under the *D. melanogaster ubiquitin* promoter (pUBI) and integrated into *attP40* sites on chromosome 2 (Stock 13-20). Extended Nxf3-GFP-3xFLAG and annotated Nxf3-GFP-3xFLAG were cloned in an in-house generated transgenesis vector for phiC31-mediated integration and expressed under the *D. melanogaster act5c* promoter (pAct5c) and integrated into *attP40* sites on chromosome 3 (Stock 13-18). Microinjection and fly stock generation was carried out by the University of Cambridge Department of Genetics Fly Facility. Mutant flies were identified by genotyping PCRs and confirmed by Sanger sequencing.

### RNA isolation and qPCR analysis

Samples were lysed in 1 ml Trizol and RNA was extracted according to manufacturer’s instructions. 1 µg of total RNA was treated with DNaseI (Thermo Fisher Scientific), and reverse transcribed with the Superscript III First Strand Synthesis Kit (Thermo Fisher Scientific), using oligo(dT)_20_ primers. Real-time PCR (qPCR) experiments were performed with a QuantStudio Real-Time PCR Light Cycler (Thermo Fisher Scientific). Transposon levels were quantified using the ΔΔCT method (Livak and Schmittgen 2001), normalized to *rp49* and fold changes were calculated relative to the indicated controls. All oligonucleotide sequences are given in **Table S2**.

### Small RNA-seq library preparation

Small RNA libraries were generated as described previously with slight modifications (McGinn and Czech 2014). Briefly, 18-to 29-nt sized small RNAs were purified by PAGE from 12 µg of total RNA from ovaries. Next, the 3’ adapter (containing 4 random nucleotides at the 5’ end (Jayaprakash et al. 2011) was ligated using T4 RNA ligase 2, truncated KQ (NEB). Following recovery of the products by PAGE purification, the 5’ adapter (containing 4 random nucleotides at the 3’ end) was ligated to the small RNAs using T4 RNA ligase (Ambion). Small RNAs containing both adapters were recovered by PAGE purification, reverse transcribed and PCR amplified. Libraries were sequenced on an Illumina HiSeq 4000.

### RNA-seq library preparation

5 µg of RNA of was cleaned up using using RNeasy mini prep column (Qiagen), according to manufacturer’s instructions. 1 µg of input RNA was used for rRNA removal with Ribo-Zero rRNA Removal Kit (Human/Mouse/Rat) (Illumina). Libraries were generated with the NEBNext Ultra Directional RNA Library Prep kit for Illumina (NEB) according to manufacturer’s instructions. The pooled libraries were quantified with KAPA Library Quantification Kit for Illumina (Kapa Biosystems) and sequenced on an Illumina HiSeq 4000 (Illumina).

### RIP-seq library preparation

RIP-seq was adapted from (D et al. 2016). Ovaries from ∼100 *w*^*1118*^ or *nxf3*^*mut*^ flies (3-5 days old) were dissected in ice-cold PBS and fixed with 0.1% PFA for 20 min, followed by quenching with equal volumes of 125 mM Glycine. Fixed ovaries were lysed in 1 ml of RIPA Buffer (supplemented with complete protease inhibitors (Roche) and RNasin Plus 40U/ml) and homogenized using a motorized pestle. Lysates were incubated 20 min at 4 °C on a tube rotator and sonicated with a Bioruptor Pico (3 cycles of 30 sec on/30sec off). After lysis, lysates were spun at 4 °C max speed for 10 min. Lysates were diluted by adding equal volume of RIP binding/wash buffer (150 mM KCl, 25 mM Tris (pH 7.5), 5 mM EDTA, 0.5 % NP-40, 0.5 mM DTT, supplemented with protease inhibitors and RNasin Plus 1:1000). Lysates were pre-cleared using 40 μl of Pierce Protein A/G beads for 1 hr at 4 °C and Nxf3 proteins were immunoprecipitated by incubation with 40 μl Nxf3 antibody overnight at 4 °C. 80 µl of Pierce A/G magnetic beads were the added to the lysates and incubated for 3 hrs at 4 °C. An aliquot of pre-cleared input lysate was saved for RNA isolation and library preparation. Following 3 washes in of RIP binding/wash buffer, IP and input samples were reverse crosslinked in 1x Reverse Crosslinking buffer (PBS, 2% N-lauroyl sarcosine, 10 mM EDTA, 5 mM DTT) and Proteinase K. RNA isolation was performed using Trizol and 50 ng of input or IP RNA were used for rRNA removal using RiboGone - Mammalian (Clontech) and library preparation using the SMARTer stranded RNA-seq Kit (Clontech). DNA libraries were quantified with KAPA Library Quantification Kit for Illumina (Kapa Biosystems) and deep-sequenced with Illumina HiSeq 4000 (Illumina).

### Ovary immunostaining

Fly ovaries were dissected in ice-cold PBS, fixed for 14 min in 4% PFA at RT and permeabilized with 3×10min washes in PBS with 0.3% Triton (PBS-Tr). Samples were blocked in PBS-Tr with 1% BSA for 2 hrs at RT and incubated overnight at 4 °C with primary antibodies in PBS-Tr and 1% BSA. After 3×10 min washes at RT in PBS-Tr, secondary antibodies were incubated overnight at 4 °C in PBS-Tr and 1% BSA. After 4×10min washes in PBS-Tr at RT (DAPI was added during the third wash) and 2×5 min washes in PBS, samples were mounted with ProLong Diamond Antifade Mountant (Thermo Fisher Scientific #P36961) and imaged on a Leica SP8 confocal microscope. Images were deconvoluted using Huygens Professional. The following antibodies were used: anti-GFP (ab13970), anti-Nxf3, anti-CG13741/Bootlegger, anti-Piwi (Brennecke et al. 2007), anti-Aub (Senti et al. 2015), anti-Ago3 (Senti et al. 2015), anti-Vasa (DSHB Cat# anti-vasa,), anti-Lamin (DSHB ADL67.10), anti-UAP56 (Eberl et al. 1997), anti-Nxt1 (Herold et al. 2001).

### Ovary immunostaining/RNA Fluorescent In Situ Hybridization (FISH)

Fly ovaries were dissected in ice-cold PBS, fixed for 14 min in 4% PFA at RT and permeabilized with 3×10min washes in PBS with 0.3% Triton (PBS-Tr). Samples were blocked in PBS-Tr with 1% BSA (supplemented with RNasin 1:1,000) for 2 hrs at RT and incubated overnight at 4 °C with primary antibodies in PBS-Tr and 1% BSA (supplemented with RNasin 1:1,000). After 3×10 min washes at RT in PBS-Tr, secondary antibodies were incubated overnight at 4 °C in PBS-Tr and 1% BSA (supplemented with RNasin 1:1,000). After 4×10 min washes in PBS-Tr, ovaries were fixed in Fixation solution (4% PFA, 0.15% Triton in PBS) for 20 min. Ovaries were washed 3×10min in PBS-Tr and permeabilized overnight in 70% ethanol. Ovaries were rehydrated in 2x SSC for 5 min. For cluster *42AB* RNA FISH, samples were pre-hybridized in 500 µl of 30% hybridization buffer (Molecular Instrument) for 30 min at 37 °C. Ovaries were resuspended in 30% hybridization buffer (Molecular Instrument) containing 1 µl of 2 µM of each probe set (even and odd) and incubated for 24 hrs at 37 °C. Samples were washed 4×15 min with 500 µl of 30% probe wash buffer (Molecular Instrument) followed by 3×5 min washes with 5x SSCT (0.1% Tween-20 in 5x SSC) at RT. Samples were pre-amplified in 500 µl amplification buffer (Molecular Instrument) for 30 min at RT. 30 pmol fluorescently labelled hairpin were heated at 95 °C for 90 sec and cooled to room temperature for 30 min. Amplification solution was prepared by adding the fluorescently labelled hairpins to 500 µl of amplification buffer. Samples were incubated overnight at RT in 125 µl of amplification solution. Samples were washed 2×5min (DAPI was added 1:5,000 to the second wash), 2×30 min, 1×5 min with 5x SSCT. Samples were mounted with ProLong Diamond Antifade Mountant (Thermo Fisher Scientific #P36961) and imaged on a Leica SP8 confocal microscope. Images were deconvoluted using Huygens Professional. The following antibodies were used: anti-Nxf3, anti-CG13741, anti-Lamin (DSHB ADL67.10).

### Image analysis

Image analysis was carried out on Fiji. For the display of germ cells nuclei, the nuclear outline was derived either from DAPI or Lamin stain. Quantification of RNA-FISH was performed by counting nuclear and cytoplasmic foci with Fiji after size and intensity thresholding, the nuclear outline was derived either from DAPI staining.

### Co-Immunoprecipitation from ovaries

Ovaries from 50 Bootlegger-GFP and 50 control flies were dissected in ice-cold PBS and lysed in 200 μl of CoIP Lysis Buffer (20 mM Tris-HCl pH 7.5, 150 mM NaCl, 2 mM MgCl2, 10% glycerol, 1 mM DTT, 0.1 mM PMSF, 0.2% NP-40 supplemented with complete protease inhibitors) and homogenized using a motorized pestle. Lysates were cleared for 5 min at 16000g and the residual pellet re-extracted with the same procedure. GFP-tagged proteins were immunoprecipitated by incubation with 25 µl of GFP-Trap magnetic agarose beads (Chromotek) overnight at 4 °C on a tube rotator. Nxf3 proteins were immunoprecipitated by incubation with 10 µl of anti-Nxf3 antibody overnight at 4 °C on a tube rotator. 10 µl of Pierce A/G magnetic beads were the added to the lysates and incubated for 3 hrs at 4 °C. The beads were washed 6x with Lysis Buffer. Samples were eluted using 20 µl of 2x sample buffer (Invitrogen) by boiling beads for 3 min and analyzed by western blot.

### Western blot

Protein concentration was measured using a Direct Detect Infrared Spectrometer (Merck). 20 µg of proteins were separated on a NuPAGE 4-12% Bis-Tris gel (Thermo Fisher Scientific). Proteins were transferred with an iBLot2 device (Invitrogen) on a nitrocellulose membrane and blocked for 1 hr in 1x Licor TBS Blocking Buffer (Licor). Primary antibodies were incubated over night at 4 °C. Licor secondary antibodies were incubated for 45 min at room temperature and images acquired with an Odyssey CLx scanner (Licor) using secondary antibodies conjugated to infra-red dyes from LiCor. The following primary antibodies were used: anti-CG13741, anti-Nxf3, anti-Tubulin (ab18251), anti-GFP (ab13790), anti-FLAG (F1804, Sigma), anti-HA (3724S, Cell signalling).

### Nxf3 pull-down and Mass Spectrometry

Ovaries from ∼100 control flies (*w*^*1118*^ strain, 3-5 days old) were dissected in ice-cold PBS and lysed in 400 μl of RIPA Buffer (supplemented with complete protease inhibitors) and homogenized using a motorized pestle. Lysates were incubated 20 min at 4 °C on a tube rotator and sonicated with a Bioruptor. Pico (3 cycles of 30 sec on/30sec off). After lysis, lysates were spun at 4 °C max speed for 10 minutes. Lysates were pre-cleared using 40 μl of Pierce Protein A/G beads for 1 hr at 4 °C and Nxf3 proteins were immunoprecipitated by incubation with 40 μl Nxf3 antibody overnight at 4 °C. 80 µl of Pierce A/G magnetic beads were the added to the lysates and incubated for 3 hrs at 4 °C. Beads were washed 3×10 min with wash buffer (150 mM KCl, 25 mM Tris (pH 7.5), 5 mM EDTA, 0.5 % NP-40, 0.5 mM DTT supplemented with complete protease inhibitors). Beads were rinsed twice with 100 mM Ammonium Bicarbonate and submitted for Mass Spectrometry. Samples were analyzed on a Q-Exactive HF mass spectrometer (Thermo Fisher Scientific) after Trypsin digestion.

Spectral .raw files were processed with the SequestHT search engine on Thermo ScientificTM Proteome Discoverer™ 2.2. Data was searched first against a custom FlyBase database (“dmel-all-translation-r6.24”) at a 1% spectrum level FDR criteria using Percolator (University of Washington). Data was also searched against a custom database including only the N-terminal extended version of Nxf3 (amino acid sequence available as **Table S3**). MS1 mass tolerance was constrained to 20 ppm and the fragment ion mass tolerance was set to 0.02 Da. Oxidation of methionine residues (+15.995 Da) AND deamidation (+0.984) of asparagine and glutamine residues were included as dynamic modifications. The Precursor Ion Quantifier node (Minora Feature Detector) included a Minimum Trace Length of 5, Max. ΔRT of Isotope Pattern 0.2 minutes. For calculation of Precursor ion intensities, Feature mapper was set True for RT alignment (mass tolerance of 10 ppm). Precursor abundance was quantified based on intensity and the level of confidence for peptide identifications was estimated using the Percolator node with a Strict FDR at q-value < 0.01.

### Cell culture

*Drosophila* Schneider 2 (S2) cells were purchased from Thermo Fisher Scientific and were grown at 26 °C in Schneider media supplemented with 10% FBS. S2 cells were transfected using Effectene (Qiagen), according to manufacturer’s instructions. S2 cells were treated with Leptomycin B (LMB, final concentration 40 ng/ml) for 12 hrs.

### S2 cells immunostaining

Cells were plated on Concanavalin A coated coverslips, fixed for 15 min in 4% PFA, permeabilized for 10 min in PBS with 0.2% Triton (PBST) and blocked for 30 min in PBS, 0.1% Tween-20 and 1% BSA. Primary antibodies were diluted in PBS, 0.1% Tween-20 and 0.1% BSA and incubated overnight at 4 °C. After 3×5 min washes in PBST, secondary antibodies were incubated for 1 hr at RT. After 3×5 min washes in PBST, DAPI was incubated for 10 min at RT, washed 2 times and the coverslips were mounted using ProLong Diamond Antifade Mountant (Thermo Fisher Scientific #P36961) and imaged on a Leica SP8 confocal microscope (100x Oil objective). The following antibodies were used: anti-Lamin (DSHB ADL67.10), anti-FLAG (14793, Cell signalling).

### Co-immunoprecipitation from cell lysates

S2 cells were transfected with 3xFLAG- and HA-tagged constructs. Cells were harvested 48 hrs after transfection and lysed in 250 μl of CoIP Lysis Buffer (Pierce) supplemented with Complete protease inhibitors (Roche). Protein lysates were diluted to 1 ml with CoIP Lysis Buffer and the 3xFLAG-tagged bait was immunoprecipitated by incubation with 20 μl of anti-FLAG M2 Magnetic Beads (Sigma M8823) for 2 hrs at 4 °C on a tube rotator. The beads were washed 3×15 min with TBS supplemented with protease inhibitors. Beads were then resuspended in 2x NuPAGE LDS Sample Buffer (Thermo Fisher Scientific) without reducing agent and boiled for 3 min at 90 °C to elute immunoprecipitated proteins. IPs, unbound fractions and input fractions were diluted to 1x NuPAGE LDS Sample Buffer concentration and reducing agent was added. Samples were boiled at 70 °C for 10 min before separating proteins by western blot.

### RNA-seq and RIP-seq analysis

Raw fastq files generated by Illumina sequencing were analyzed by a pipeline developed in-house. In short, the 5 first bases and the last base of each 50 bp read were removed using fastx trimmer (http://hannonlab.cshl.edu/fastx_toolkit/). Following removal of rRNA mapped reads, high-quality reads were aligned to transposon consensus sequences followed by alignment to the *Drosophila melanogaster* genome release 6 (dm6; downloaded from Flybase) using STAR (Dobin et al. 2013). Genome multi-mapping reads were randomly assigned to one location using option ‘--outFilterMultimapNmax 1000 --outMultimapperOrder Random’ and non-mapping reads were removed. For genome-wide analyses, unique mapper reads were extracted to ensure unique locations of reads. Normalization was achieved by calculating rpm (reads per million). Reads mapping to genes were counted with htseq (Anders et al. 2015) and transposon derived reads were calculated using a custom script. Differential expression analysis was performed using custom built R scripts. piRNA clusters were divided in 1 kb bin and unique reads mapping to these bins were counted using htseq. A pseudo count of 0.01 or 0.1 respectively for RNA-seq and RIP-seq experiment, was then added to each bin before calculating log_2_ fold-changes per bin.

### small RNA-seq analysis

For small RNA-seq, adapters were clipped from raw fastq files with fastx_clipper (adapter sequence AGATCGGAAGAGCACACGTCTGAACTCCAGTCA) keeping only reads with at least 23 bp length. Then the first and last 4 bases were trimmed using seqtk (https://github.com/lh3/seqtk). After removal of cloning markers and 2S rRNA mapped reads, alignment was performed as described above and normalized to miRNA reads in the control library (set to rpm). Only high-quality small RNA reads with a length between 18 and 30 bp were used for further analysis of small RNA profiles. piRNA distribution was calculated and plotted in R. piRNA clusters were divided in 1 kb bin and unique reads mapping to these bins were counted using htseq. A pseudo count of 1 was added to each bin before calculating log_2_ fold-changes per bin. Small RNA size profiles were plotted in R, only unique mappers were used for small RNA distribution of piRNA clusters.

### Data availability

Sequencing data reported in this paper has been deposited in Gene Expression Omnibus (GSE133528). Mass Spectrometry data has been deposited to the PRIDE Archive (PXD014472).

## References

Akkouche A, Mugat B, Barckmann B, Varela-Chavez C, Li B, Raffel R, Pelisson A, Chambeyron S. 2017. Piwi Is Required during Drosophila Embryogenesis to License Dual-Strand piRNA Clusters for Transposon Repression in Adult Ovaries. Mol Cell 66: 411–419 e414.

Anders S, Pyl PT, Huber W. 2015. HTSeq--a Python framework to work with high-throughput sequencing data. Bioinformatics 31: 166–169.

Andersen PR, Tirian L, Vunjak M, Brennecke J. 2017. A heterochromatin-dependent transcription machinery drives piRNA expression. Nature 549: 54–59.

Batki J, Schnabl J, Wang J, Handler D, Andreev VI, Stieger CE, Novatchkova M, Lampersberger L, Kauneckaite K, Xie W et al. 2019. The nascent RNA binding complex SFiNX licenses piRNA-guided heterochromatin formation. bioRxiv doi: https://doi.org/10.1101/609693.

Bjork P, Wieslander L. 2017. Integration of mRNP formation and export. Cell Mol Life Sci 74: 2875–2897.

Black BE, Holaska JM, Levesque L, Ossareh-Nazari B, Gwizdek C, Dargemont C, Paschal BM. 2001. NXT1 is necessary for the terminal step of Crm1-mediated nuclear export. J Cell Biol 152: 141–155.

Brennecke J, Aravin AA, Stark A, Dus M, Kellis M, Sachidanandam R, Hannon GJ. 2007. Discrete small RNA-generating loci as master regulators of transposon activity in Drosophila. Cell 128: 1089–1103.

Chen YA, Stuwe E, Luo Y, Ninova M, Le Thomas A, Rozhavskaya E, Li S, Vempati S, Laver JD, Patel DJ et al. 2016. Cutoff Suppresses RNA Polymerase II Termination to Ensure Expression of piRNA Precursors. Mol Cell 63: 97–109.

Cheng H, Dufu K, Lee CS, Hsu JL, Dias A, Reed R. 2006. Human mRNA export machinery recruited to the 5’ end of mRNA. Cell 127: 1389–1400.

Czech B, Munafo M, Ciabrelli F, Eastwood EL, Fabry MH, Kneuss E, Hannon GJ. 2018. piRNA-Guided Genome Defense: From Biogenesis to Silencing. Annu Rev Genet 52: 131–157.

Czech B, Preall JB, McGinn J, Hannon GJ. 2013. A transcriptome-wide RNAi screen in the Drosophila ovary reveals factors of the germline piRNA pathway. Mol Cell 50: 749–761.

D GH Kelley DR, Tenen D, Bernstein B, Rinn JL. 2016. Widespread RNA binding by chromatin-associated proteins. Genome Biol 17: 28.

Dennis C, Brasset E, Sarkar A, Vaury C. 2016. Export of piRNA precursors by EJC triggers assembly of cytoplasmic Yb-body in Drosophila. Nat Commun 7: 13739.

Dobin A, Davis CA, Schlesinger F, Drenkow J, Zaleski C, Jha S, Batut P, Chaisson M, Gingeras TR. 2013. STAR: ultrafast universal RNA-seq aligner. Bioinformatics 29: 15–21.

Eberl DF, Lorenz LJ, Melnick MB, Sood V, Lasko P, Perrimon N. 1997. A new enhancer of position-effect variegation in Drosophila melanogaster encodes a putative RNA helicase that binds chromosomes and is regulated by the cell cycle. Genetics 146: 951–963.

ElMaghraby MF, Andersen PR, Pühringer F, Meixner K, Lendl T, Tirian L, Brennecke J. 2019. A heterochromatin-specific RNA export pathway facilitates piRNA production. bioRxiv doi: https://doi.org/10.1101/596171.

Fabry MH, Ciabrelli F, Munafo M, Eastwood EL, Kneuss E, Falciatori I, Falconio FA, Hannon GJ, Czech B. 2019. piRNA-guided co-transcriptional silencing coopts nuclear export factors. Elife 8.

Goriaux C, Desset S, Renaud Y, Vaury C, Brasset E. 2014. Transcriptional properties and splicing of the flamenco piRNA cluster. EMBO Rep 15: 411–418.

Handler D, Meixner K, Pizka M, Lauss K, Schmied C, Gruber FS, Brennecke J. 2013. The genetic makeup of the Drosophila piRNA pathway. Mol Cell 50: 762–777.

Herold A, Klymenko T, Izaurralde E. 2001. NXF1/p15 heterodimers are essential for mRNA nuclear export in Drosophila. RNA 7: 1768–1780.

Jayaprakash AD, Jabado O, Brown BD, Sachidanandam R. 2011. Identification and remediation of biases in the activity of RNA ligases in small-RNA deep sequencing. Nucleic Acids Res 39: e141.

Klattenhoff C, Xi H, Li C, Lee S, Xu J, Khurana JS, Zhang F, Schultz N, Koppetsch BS, Nowosielska A et al. 2009. The Drosophila HP1 homolog Rhino is required for transposon silencing and piRNA production by dual-strand clusters. Cell 138: 1137–1149.

Kohler A, Hurt E. 2007. Exporting RNA from the nucleus to the cytoplasm. Nat Rev Mol Cell Biol 8: 761–773.

Kudo N, Matsumori N, Taoka H, Fujiwara D, Schreiner EP, Wolff B, Yoshida M, Horinouchi S. 1999. Leptomycin B inactivates CRM1/exportin 1 by covalent modification at a cysteine residue in the central conserved region. Proc Natl Acad Sci U S A 96: 9112–9117.

Le Hir H, Gatfield D, Izaurralde E, Moore MJ. 2001. The exon-exon junction complex provides a binding platform for factors involved in mRNA export and nonsense-mediated mRNA decay. EMBO J 20: 4987–4997.

Lim AK, Kai T. 2007. Unique germ-line organelle, nuage, functions to repress selfish genetic elements in Drosophila melanogaster. Proc Natl Acad Sci U S A 104: 6714–6719.

Livak KJ, Schmittgen TD. 2001. Analysis of relative gene expression data using real-time quantitative PCR and the 2(-Delta Delta C(T)) Method. Methods 25: 402–408.

Malone CD, Brennecke J, Dus M, Stark A, McCombie WR, Sachidanandam R, Hannon GJ. 2009. Specialized piRNA pathways act in germline and somatic tissues of the Drosophila ovary. Cell 137: 522–535.

McGinn J, Czech B. 2014. Small RNA library construction for high-throughput sequencing. Methods Mol Biol 1093: 195–208.

Mohn F, Sienski G, Handler D, Brennecke J. 2014. The rhino-deadlock-cutoff complex licenses noncanonical transcription of dual-strand piRNA clusters in Drosophila. Cell 157: 1364–1379.

Muerdter F, Guzzardo PM, Gillis J, Luo Y, Yu Y, Chen C, Fekete R, Hannon GJ. 2013. A genome-wide RNAi screen draws a genetic framework for transposon control and primary piRNA biogenesis in Drosophila. Mol Cell 50: 736–748.

Murano K, Iwasaki YW, Ishizu H, Mashiko A, Shibuya A, Kondo S, Adachi S, Suzuki S, Saito K, Natsume T et al. 2019. Nuclear RNA export factor variant initiates piRNA-guided cotranscriptional silencing. bioRxiv doi: https://doi.org/10.1101/605725.

Ohno M, Segref A, Bachi A, Wilm M, Mattaj IW. 2000. PHAX, a mediator of U snRNA nuclear export whose activity is regulated by phosphorylation. Cell 101: 187–198.

Ossareh-Nazari B, Maison C, Black BE, Levesque L, Paschal BM, Dargemont C. 2000. RanGTP-binding protein NXT1 facilitates nuclear export of different classes of RNA in vitro. Mol Cell Biol 20: 4562–4571.

Ozata DM, Gainetdinov I, Zoch A, O’Carroll D, Zamore PD. 2019. PIWI-interacting RNAs: small RNAs with big functions. Nat Rev Genet 20: 89–108.

Pane A, Jiang P, Zhao DY, Singh M, Schupbach T. 2011. The Cutoff protein regulates piRNA cluster expression and piRNA production in the Drosophila germline. EMBO J 30: 4601–4615.

Port F, Chen HM, Lee T, Bullock SL. 2014. Optimized CRISPR/Cas tools for efficient germline and somatic genome engineering in Drosophila. Proc Natl Acad Sci U S A 111: E2967–2976.

Senti KA, Jurczak D, Sachidanandam R, Brennecke J. 2015. piRNA-guided slicing of transposon transcripts enforces their transcriptional silencing via specifying the nuclear piRNA repertoire. Genes Dev 29: 1747–1762.

Stewart M. 2010. Nuclear export of mRNA. Trends Biochem Sci 35: 609–617.

Yang J, Bogerd HP, Wang PJ, Page DC, Cullen BR. 2001. Two closely related human nuclear export factors utilize entirely distinct export pathways. Mol Cell 8: 397–406.

Yi R, Qin Y, Macara IG, Cullen BR. 2003. Exportin-5 mediates the nuclear export of pre-microRNAs and short hairpin RNAs. Genes Dev 17: 3011–3016.

Yoh SM, Cho H, Pickle L, Evans RM, Jones KA. 2007. The Spt6 SH2 domain binds Ser2-P RNAPII to direct Iws1-dependent mRNA splicing and export. Genes Dev 21: 160–174.

Zhang F, Wang J, Xu J, Zhang Z, Koppetsch BS, Schultz N, Vreven T, Meignin C, Davis I, Zamore PD et al. 2012. UAP56 couples piRNA clusters to the perinuclear transposon silencing machinery. Cell 151: 871–884.

Zhang G, Tu S, Yu T, Zhang XO, Parhad SS, Weng Z, Theurkauf WE. 2018. Co-dependent Assembly of Drosophila piRNA Precursor Complexes and piRNA Cluster Heterochromatin. Cell Rep 24: 3413–3422 e3414.

Zhang Z, Wang J, Schultz N, Zhang F, Parhad SS, Tu S, Vreven T, Zamore PD, Weng Z, Theurkauf WE. 2014. The HP1 homolog rhino anchors a nuclear complex that suppresses piRNA precursor splicing. Cell 157: 1353–1363.

Zhao K, Cheng S, Miao N, Xu P, Lu X, Zhang Y, Wang M, Ouyang X, Yuan X, Liu W et al. 2019. A Pandas complex adapted for piRNA-guided transposon silencing. bioRxiv doi: https://doi.org/10.1101/608273.

